# Cohesin and NuRD Antagonistically Drive Alternative Neuronal Fates via PLZF Transcription Factors

**DOI:** 10.64898/2026.04.16.718828

**Authors:** Dongyeop Lee, Takashi Hirose, H. Robert Horvitz

## Abstract

Diverse genetic and epigenetic factors cooperate to specify cellular fates during development. Establishing these fates is especially critical in the nervous system, which comprises diverse neuronal cell types. How genomic architecture interfaces with epigenetic regulators to drive transcriptional programs underlying neuronal fates remains poorly understood. Here we show that cohesin, a protein complex that shapes genomic architecture, promotes GABAergic fate specification in a subset of neurons in the nematode *Caenorhabditis elegans*. This process is facilitated by EOR-1, a homolog of the human promyelocytic leukemia zinc finger (PLZF) transcription factor. The nucleosome remodeling and deacetylase (NuRD) complex and TRA-4, another PLZF homolog, promote tyraminergic fate in the normally GABAergic neurons when cohesin or EOR-1 function is lost, revealing an antagonistic mechanism determining alternative neuronal fates. These findings highlight a critical interplay among genome architecture, epigenetic remodeling and transcriptional regulation in neuronal fate specification and, given the evolutionary conservation of these factors, suggest a mechanism underlying neural development across species.

**Teaser:** Genomic architecture, epigenetic and genetic factors interact to determine specific neuronal fates during development.

## Introduction

The evolutionarily conserved cohesin protein complex plays a critical role in modulating DNA architecture. The subunits of cohesin form a ring-like structure that alters DNA conformation to influence diverse cellular processes, including sister chromatid cohesion, DNA repair and replication (*1*). In humans, mutations in cohesin genes are associated with genetic developmental and neurodevelopmental disorders that present with severe multisystem abnormalities, such as growth retardation, microcephaly and cognitive disabilities (*2–8*). Alterations in genomic structure have been linked to chromatin remodeling and epigenetic modifications, which in turn regulate the recruitment of transcription factors to specific genomic regions to effect changes in gene expression (*9*). During development, the fate of an individual cell is determined by interactions among specific genetic and epigenetic factors at precise developmental stages, and the expression of specific genes in specific cells at specific times is essential for the proper determination of neuronal cell fates (*10*). Nonetheless, how cohesin influences cellular development and what factors interact with cohesin during development remain unclear.

Because of its simple and completely known neuroanatomy, completely known cell lineage, and transparent body, *Caenorhabditis elegans* is an excellent organism for analyses of the molecular genetic mechanisms that generate neuronal diversity. The nervous system of the *C. elegans* hermaphrodite contains 302 neurons of 118 classes with distinct anatomies, connectivities and neurotransmitters (e.g., acetylcholine, GABA, glutamate, dopamine, serotonin, octopamine and tyramine) (*11*). Much is known about genes encoding transcription factors that contribute to this neuronal diversity (*12*). However, how transcription factors, epigenetic factors and genomic architecture interact in specifying neuronal cell fates remains unexplored.

Here we report that cohesin determines neuronal cell identity by interacting with a PLZF (promyelocytic leukemia zinc finger) transcription factor. Specifically, we found that cohesin and EOR-1, a *C. elegans* ortholog of human PLZF, function together to drive a specific GABAergic neuronal fate instead of an alternative tyraminergic fate. Conversely, the NuRD (nucleosome remodeling and deacetylase) complex and TRA-4, another *C. elegans* PLZF ortholog, drive the tyraminergic fate and hence function antagonistically to cohesion and EOR-1. These observations reveal a functional interaction among cohesin, PLZF-like transcription factors, and chromatin factors in the generation of specific neuronal identities.

## Results

### Cohesin mutants generate extra adrenergic-like cells in developing *C. elegans*

*C. elegans* has two pairs of adrenergic neurons – two tyraminergic RIML/R (RIM Left and RIM Right) and two octopaminergic RICL/R (RIC Left and RIC Right) neurons – all of which are located in the head (**Fig. 1A**). Tyramine is synthesized from tyrosine by the enzymatic activity of tyrosine decarboxylase, which is encoded by the gene *tdc-1*. In octopaminergic cells, tyramine is further processed by tyramine beta-hydroxylase (encoded by the gene *tbh-1*) to produce octopamine. To identify genes that regulate the development of adrenergic neurons, we expressed the green fluorescence protein (GFP) with a nuclear-localization signal (NLS) under the control of the *tdc-1* promoter (*tdc-1p::4xNLS::GFP*), thereby driving GFP expression specifically in the nuclei of the four adrenergic neurons (the two RIMs and the two RICs) (**Fig. 1A**). We performed ethyl methanesulfonate (EMS) mutagenesis screens for mutants with extra or fewer GFP-positive cells in the head and identified the mutant genes responsible using whole-genome sequencing (**Fig. 1B**). Among the mutations that caused a reduced number of GFP-positive cells, we identified an allele of the gene *hlh-13*, which was reported to be a regulator of RIC cell fates while we were preparing our results for publication (*13*). Another isolate, which carried the mutation *n6608*, displayed extra GFP-positive cells in the head and proved to have a premature stop codon (opal) in the last exon of the gene *coh-1* (**Fig. 1C**). *coh-1* encodes a homolog of yeast and human RAD21, a subunit of the cohesin complex; the *C. elegans* genome contains five paralogs of RAD21 (**Fig. 1D**). To confirm that the *coh-1(n6608)* mutation was responsible for the generation of the extra GFP-positive cells, we used CRISPR/Cas9 to generate *coh-1(n6618)*, which contains a mutation identical to that of *n6608* (**Fig. 1C**). *coh-1(n6618)* mutants also showed extra GFP-positive cells, and the penetrance was comparable to that of *coh-1(n6608)* (**Fig. 1, E** and **F**). Expression of the wild-type *coh-1* gene fused with wrmScarlet (*coh-1::wrmScarlet*) rescued the generation of extra adrenergic-like cells in *coh-1(n6608)* mutants, suggesting that a loss of *coh-1* function is responsible for the phenotype (**Fig. 1G**). To test whether *coh-1(n6608)* and *coh-1(n6618)* alleles are null, we created a knockout *coh-1* allele that deletes the entire coding region. Homozygous *coh-1* knockout mutants arrested and died as larvae, indicating that *coh-1(n6608)* and *coh-1(n6618)* are reduction-of-function and not null alleles. These results also suggest that *coh-1* is essential for viability and that its function cannot be compensated by other COH-1 paralogs. Cohesin has two distinct functions, a cohesive function (which acts in sister chromatid exchange) and an extrusive function (which acts in DNA loop extrusion) (*13*) (**Fig. 1H**). Because *C. elegans* COH-1 is not involved in mitosis (*14*), a process that requires sister chromatid cohesion, we suggest that it is cohesin’s role in DNA loop extrusion that is critical for preventing the generation of extra adrenergic-like cells.

**Fig. 1.**
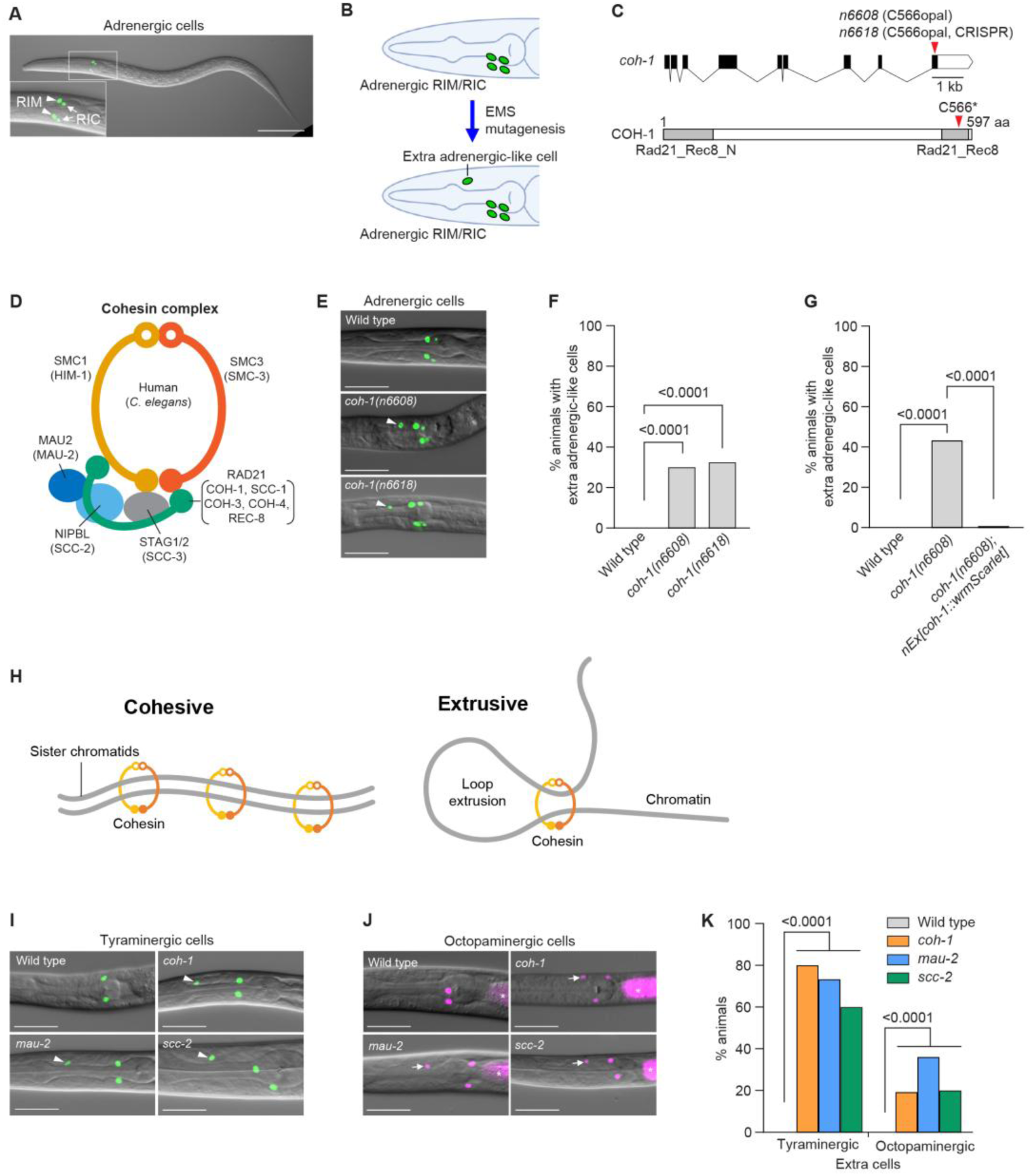
*coh-1* cohesin mutants generate extra tyraminergic-like and octopaminergic-like cells. (**A**) The *tdc-1p::4xNLS::GFP* (adrenergic reporter) is expressed in the two lateral pairs of tyraminergic (RIML and RIMR) and octopaminergic (RICL and RICR) neurons in an L2-stage larval hermaphrodite. Scale bar, 50 µm. (**B**) Schematic showing an EMS mutagenesis screen to identify mutants with extra adrenergic-like cells. (**C**) The *coh-1(n6608)* mutation generates a premature stop codon (opal) in the last exon of the *coh-1* gene. *coh-1(n6618)* is the identical mutation created by CRISPR/Cas9. The Rad21_Rec8_N and Rad21_Rec8 domains are highlighted in grey. aa, amino acid. (**D**) Schematic structure of the cohesin complex adapted from Shi et al. (2020) (*95*). *C. elegans* orthologs are shown in parentheses. (**E** and **F**) *tdc-1p::4xNLS::GFP* (adrenergic reporter) expression (**E**) and the percentages of animals with extra adrenergic cells (arrowheads) (**F**) in the heads of wild-type animals and *coh-1(n6608)* and *coh-1(n6618)* mutants. Scale bar, 20 µm. p-value, Fisher’s exact test. n=120 animals. Worms at the L2-L4 larval stages were analyzed for panels **E** to **G**, and **I** to **N**. (**G**) Percentages of animals displaying extra adrenergic-like cells for wild-type and *coh-1(n6608)* mutants and *coh-1(n6608)* mutants carrying an extrachromosomal array of wild-type *coh-1* fused with wrmScarlet at the 3’ end (*nEx3168[coh-1::wrmScarlet]*). p-value, Fisher’s exact test. n=120 animals. Animals expressed a *tdc-1p::4xNLS::GFP* (adrenergic) reporter. (**H**) Cartoons showing cohesive (left) and extrusive (right) cohesin. (**I** and **J**) *coh-1(n6618)*, *mau-2(qm160)* and *scc-2(n6656)* mutants generate extra tyraminergic-like (**I**) and octopaminergic-like cells (**J**). Animals expressed either an *F23H12.7p::4xNLS::GFP* (tyraminergic) or a *tbh-1p::4xNLS::mCherry* (octopaminergic) reporter in panels **I** to **K**. Arrowheads, extra tyraminergic-like cells. Arrows, extra octopaminergic-like cells. Asterisks, co-injection marker (*ges-1p::mCherry*). Scale bar, 20 µm. (**K**) Percentages of wild-type animals and *coh-1(n6618)*, *mau-2(qm160)* and *scc-2(n6656)* mutants that have extra tyraminergic-like and extra octopaminergic-like cells. p-value, Fisher’s exact test. n=150 animals.

The *tdc-1p::4xNLS::GFP* adrenergic neuronal reporter we used is expressed in both tyraminergic RIMs and octopaminergic RICs. To determine whether the extra GFP-positive cells were extra tyraminergic- and/or extra octopaminergic-like cells, we examined tyraminergic- and octopaminergic-specific mCherry reporters in *coh-1* mutants. By analyzing the published *C. elegans* Neuronal Gene Expression Network (CeNGEN) (*15*, *16*), we identified *F23H12.7*, a nematode-specific gene that is highly expressed in the tyraminergic RIM neurons but not in other neurons. To confirm this finding, we created a *F23H12.7* promoter-driven nuclear-localized mCherry reporter (*F23H12.7p::4xNLS::mCherry*) and demonstrated that *F23H12.7* is specifically expressed in the tyraminergic RIMs and – importantly – excluded from the octopaminergic RICs (**fig. S1A**). Using this tyraminergic reporter and, separately, the octopaminergic reporter *tbh-1p::4xNLS::mCherry*, we showed that *coh-1* mutants generate both extra tyraminergic-like and extra octopaminergic-like cells (**fig. S1, A** and **B**). Additionally, the extra tyraminergic-like cells and extra octopaminergic-like cells are not the same cells, since the tyraminergic GFP and octopaminergic mCherry reporters were not co-expressed in the same cells (**fig. S1C**). To explore the possibility that the cohesin complex – as opposed to the single cohesin subunit COH-1 – functions to prevent the generation of extra tyraminergic-like and octopaminergic-like cells, we examined deletion alleles of two additional cohesin genes, the *mau-2(qm160)* null allele (*17*) and the *scc-2(n6656)* allele (the latter of which is an in-frame deletion and likely not a null allele); we generated *scc-2(n6656)* using CRISPR/Cas9 (see Materials and Methods). Similar to *coh-1* mutants, *mau-2* and *scc-2* mutants displayed both extra tyraminergic-like and extra octopaminergic-like cells (**Fig. 1, I** to **K**). Furthermore, worms treated with RNAi directed against genes encoding the three additional cohesin subunits – *smc-3*, *scc-3* and *him-1* – similarly generated extra tyraminergic-like and extra octopaminergic-like cells, although the effect of *him-1* RNAi on the generation of extra octopaminergic-like cells was not statistically significant (**fig. S1, D** to **F**). Together, these results demonstrate that disruption of the cohesin complex – not a specific *coh-1* mutation – causes the generation of extra tyraminergic-like and octopaminergic-like cells, indicating that the cohesin complex normally prevents the formation of ectopic adrenergic-like cells.

### Cohesin prevents a subset of GABAergic neurons from acquiring characteristics of an alternative tyraminergic fate

Given the stronger penetrance of the generation of extra tyraminergic-like cells than extra octopaminergic-like cells in cohesin mutants and the varied locations of extra octopaminergic-like cells (**Fig. 1K** and **fig. S1G**), we prioritized investigating the source(s) of the extra tyraminergic-like cells. To seek the origin(s) of the cells that acquired a tyraminergic-like state in cohesin mutants, we used the NeuroPAL transgenic *C. elegans* strain, which labels individual neurons with distinct sets of fluorescence colors (*18*). We generated NeuroPAL transgenic strains with the *scc-2* mutant and tyraminergic GFP reporter backgrounds (**Fig. 2A**). By analyzing the positions of the extra tyraminergic-like cell and the colors of neighboring cells, we determined that the extra tyraminergic-like cell was likely the RMED (RME Dorsal) neuron, one of the four classes of RME GABAergic motor neurons (**Fig. 2, A** and **B**). In GABAergic neurons, glutamic acid decarboxylase (GAD/UNC-25) converts glutamate to GABA, and a vesicular GABA transporter (VGAT/UNC-47) mediates GABA transport and secretion (**Fig. 2C**). To confirm our identification of the extra tyraminergic cell as RMED, we used the GABA transporter gene *unc-47* fused with histone H2B and “optimized” mCherry (*unc-47::SL2::H2B::mChOpti*) as a reporter expressed in the nuclei of GABAergic neurons (**Fig. 2D**). Indeed, our analysis showed that in cohesin mutants (*coh-1*, *mau-2* and *scc-2*), most worms (83-91%) expressed the tyraminergic reporter in RMED or in both RMED and RMEV (RME Ventral) cells, with a very small fraction of worms (3-6%) expressing the tyraminergic reporter in RMEL (RME Left) or RMER (RME Right) (**Fig. 2, D** and **E**). RNAi inhibition of genes encoding other cohesin subunits, *smc-3*, *scc-3*, or *him-1*, similarly generated tyraminergic-like RMED and RMEV cells (**fig. S2, A** and **B**). The extra tyraminergic-like cells appeared in late-stage (three-fold) embryos (**fig. S2C**), indicating that the extra tyraminergic-like cells were embryonic, as are RME neurons (*19*).

**Fig. 2.**
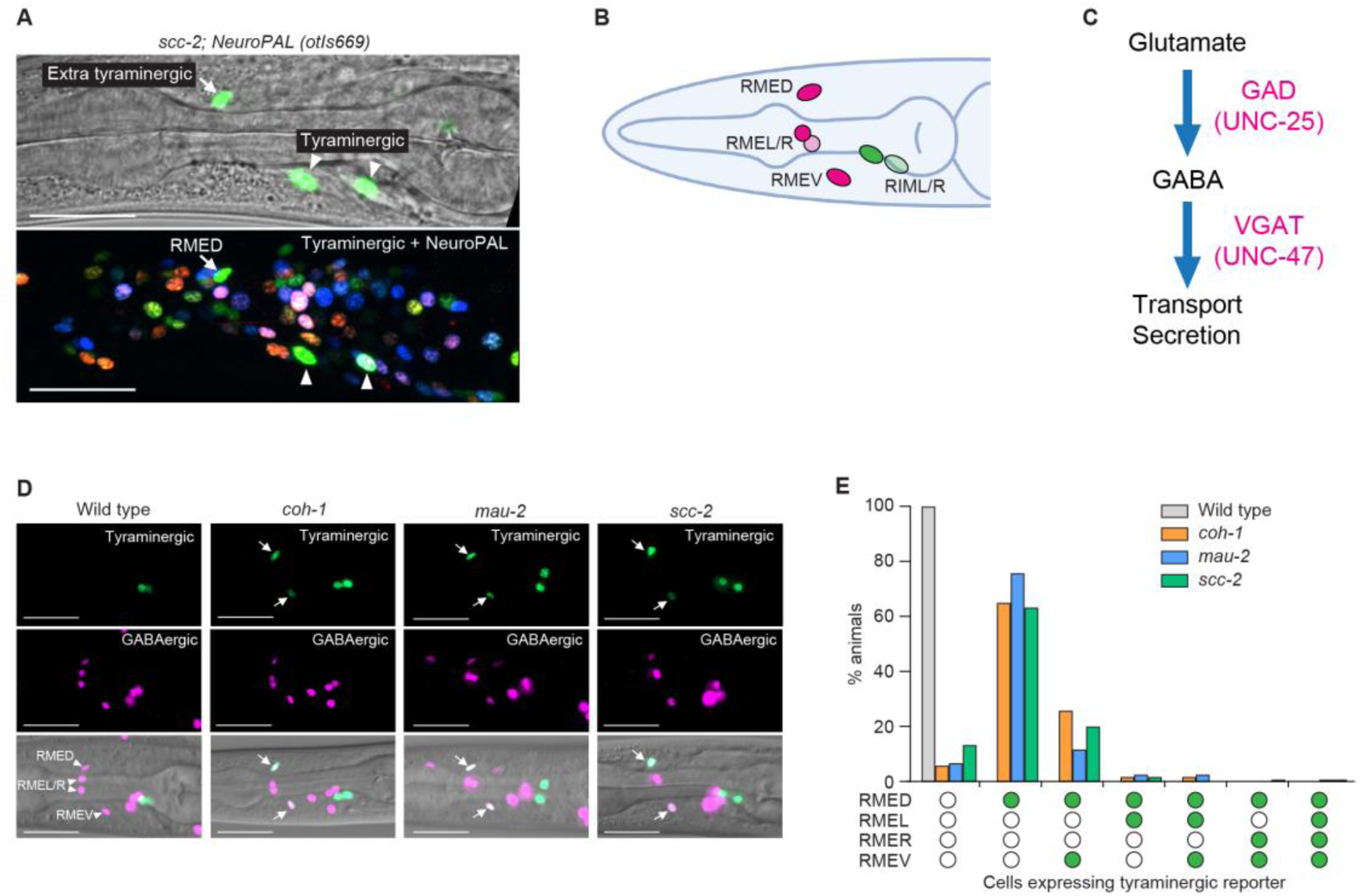
Cohesin inhibits tyraminergic fate transformation by specific GABAergic neurons. (**A**) Confocal fluorescence micrographs showing expression of *F23H12.7p::4xNLS::GFP* (tyraminergic) and NeuroPAL (*otIs669*) reporters in an *scc-2(n6656)* mutant in the day-1 adult stage. The position of the extra tyraminergic-like cell and the colors of surrounding cells identify the extra tyraminergic-like cell as RMED. The weak GFP signal detected in the pharyngeal region (upper panel) was likely generated by bleed-through or artifact from overexpression. Arrows, extra tyraminergic-like cells. Arrowheads, tyraminergic RIMs. Scale bar, 20 µm. (**B**) Schematic showing the four GABAergic RME neurons (RMED, RMEL, RMER and RMEV) and the two tyraminergic RIM neurons. (**C**) Simplified diagram showing GABA biosynthetic and transport pathways. GAD, glutamic acid decarboxylase. VGAT, vesicular GABA transporter. (**D**) The extra tyraminergic-like cells in *coh-1(n6618)*, *mau-2(qm160)* and *scc-2(n6656)* mutants are GABAergic RMED and RMEV. Animals expressed a *F23H12.7p::4xNLS::GFP* (tyraminergic) and an *unc-47::SL2::H2B::mChOpti* (GABAergic) reporter in panels **D** and **E**. Arrowheads, GABAergic RMED, RMEL, RMER and RMEV. Arrows, extra tyraminergic-like cells. Scale bar, 20 µm. Worms at the L4 larval stage were analyzed for panels **D** and **E**. (**E**) Percentages of wild-type animals and *coh-1(n6618)*, *mau-2(qm160)* and *scc-2(n6656)* mutants that have RME neurons expressing the tyraminergic-like reporter *F23H12.7p::4xNLS::GFP*. Empty circle, RME with no GFP expression. Green circle, RME with GFP expression. n=120 animals.

The *C. elegans* nervous system is known to produce seven major neurotransmitters/neuromodulators: acetylcholine, glutamate, GABA, dopamine, serotonin, tyramine and octopamine (*20–33*). To determine if the GABAergic RMED neuron that displayed a tyraminergic fate in *coh-1* mutants also acquired aspects of other neuronal fates, we examined the expression of reporters for all seven neurotransmitters in *coh-1* mutant RMED cells: octopaminergic (tyramine beta-hydroxylase, *tbh-1p::GFP*), cholinergic (choline acetyltransferase/vesicular acetylcholine transporter, *cha-1*/*unc-17p::GFP*), glutamatergic (vesicular glutamate transporter, *eat-4::SL2::YFP::H2B*), dopaminergic (dopamine transporter, *dat-1::T2A::NeonGreen*) and serotonergic (tryptophan hydroxylase, *tph-1p::GFP*) (**fig. S2, D** and **E**). Unlike the tyraminergic/octopaminergic reporter *tdc-1p::4xNLS::GFP*, which as shown above was expressed in RMED, none of the other neurotransmitter reporters we tested was expressed in RMED in *coh-1* mutants (**fig. S2, D** and **E**). Taken together, these data indicate that cohesin functions to prevent a subset of GABAergic neurons from acquiring a tyraminergic fate.

### Cohesin promotes multiple aspects of the RMED and RMEV GABAergic neuronal fate

We asked if the RMED/V neurons in cohesin mutants express GABAergic neuronal genes. Cohesin mutants expressed VGAT (*unc-47::SL2::H2B::mChOpti*) (**Fig. 2D** and **Fig. 3A**). By contrast, cohesin mutants failed to express detectable levels of GAD promoter-driven GFP (*unc-25p::GFP*) in RMED or in both RMED and RMEV, indicating that RMED/V are not GABAergic, and at least one aspect of a GABAergic fate is expressed. (**Fig. 3, A** and **B**). To further confirm this result, we sought a reporter for the RMED and RMEV neurons. We analyzed data from the *C. elegans* CeNGEN gene-expression atlas (*15*, *16*) and found that *ins-13*, which encodes an insulin-like peptide, is expressed in RMED and RMEV but not in RMEL or RMER. We generated an *ins-13* promoter-driven GFP (*ins-13p::GFP*) reporter and confirmed that *ins-13* is expressed in the RMED/Vs but not in the RMER/Ls (**Fig. 3C**). Cohesin mutants exhibited defects in *ins-13p::GFP* expression in RMED or both RMED and RMEV (**Fig. 3D**). Because 50-70% of cohesin mutants still expressed *ins-13p::GFP* in RMED and RMEV, we were able to examine the morphologies of these neurons in cohesin mutant larvae. Interestingly, the normally long processes of RMED and RMEV were truncated in cohesin mutants, and the RMED process was undetectable in the majority of cohesin mutant animals (**Fig. 3, E** to **G**). These data indicate that cohesin not only inhibits a tyraminergic fate but also promotes multiple aspects of the normal GABAergic RMED/V fate – neurotransmitter expression and axonal morphology.

**Fig. 3.**
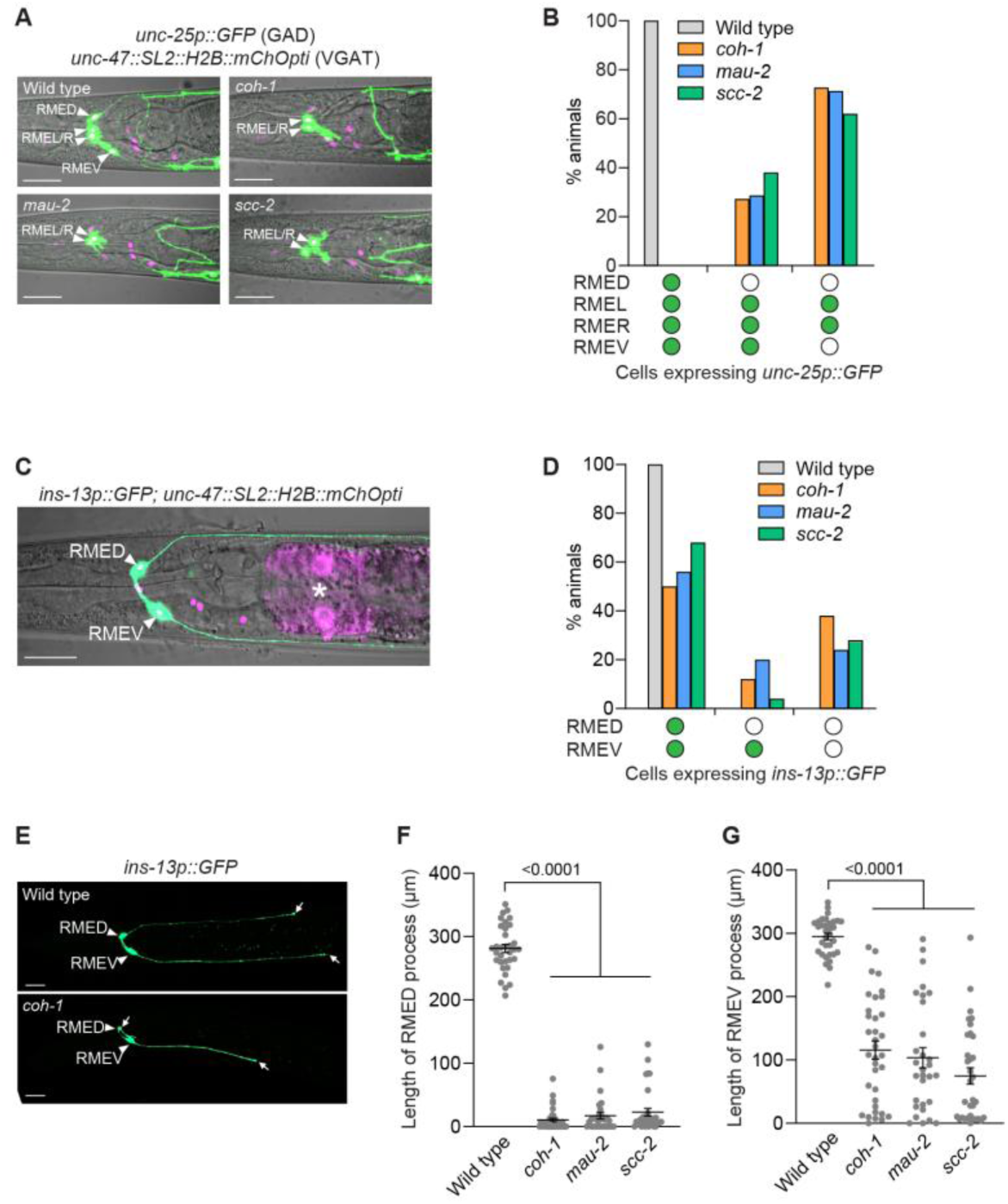
Cohesin promotes the normal development of GABAergic RMED and RMEV. (**A**) Confocal fluorescence micrographs showing that GAD is not expressed in RMED or RMEV in *coh-1(n6618)*, *mau-2(qm160)* and *scc-2(n6656)* mutants. Strains expressed *unc-25p::GFP* (GAD) and *unc-47::SL2::H2B::mChOpti* (VGAT) reporters in panels **A** and **B**. Arrowheads, RMED and RMEV. Scale bar, 20 µm. Worms at the L4 larval stage were analyzed in panels **A** to **G**. (**B**) Percentages of wild-type animals and *coh-1(n6618)*, *mau-2(qm160)* and *scc-2(n6656)* mutants that express *unc-25p::GFP* (GAD) in RMEs. Empty circle, RME with no GFP expression. Green circle, RME with GFP expression. n=150 animals. (**C**) Confocal fluorescence micrograph showing the expression of *ins-13p::GFP* in RMED and RMEV and of *unc-47::SL2::H2B::mChOpti* in an L4-larval stage of wild-type animal. Arrowheads, RMED and RMEV. Asterisk, co-injection marker (*ges-1p::mCherry*). Scale bar, 20 µm. (**D**) Percentages of wild-type animals and *coh-1(n6618)*, *mau-2(qm160)* and *scc-2(n6656)* mutants expressing *ins-13p::GFP* in RMED and/or RMEV. Empty circle, RME with no GFP expression. Green circle, RME with GFP expression. n=50 animals. (**E**) Confocal fluorescence micrographs showing *ins-13p::GFP* expression in the cell bodies and processes of RMED and RMEV in a wild-type animal and a *coh-1(n6618)* mutant. Arrowheads, RMED and RMEV. Arrows, posterior ends of RMED and RMEV processes. Scale bar, 20 µm. (**F** and **G**) Quantification of the length of RMED (**F**) and RMEV (**G**) in wild-type animals and *coh-1(n6618)*, *mau-2(qm160)* and *scc-2(n6656)* mutants. Strains expressed an *ins-13p::GFP* (RMED/V) reporter. Error bars, mean ± SEM. p-value, one-way ANOVA. n=33 animals for wild type, 35 animals for *coh-1*, 30 animals for *mau-2* and 33 animals for *scc-2(n6656)*. Please note that the same wild-type conditions were used in **fig. S4, J** and **K**.

### The known RME regulator genes *nhr-67* and *ceh-10* do not act in the cohesin pathway

Given the defects in the expression of a GABAergic reporter in RMED and RMEV in cohesin mutants, we hypothesized that genes that promote RME development might act in the cohesin pathway. We tested whether the activities of two known RME identity regulators, *nhr-67* (*32*) and *ceh-10* (*32*), contribute to the GABAergic-tyraminergic RME decision. We first asked if *nhr-67* is involved in the GABAergic-to-tyraminergic fate transformation. We expressed fluorescence reporters for RME and RIM in the background of the cold-sensitive *nhr-67(ot202*ts) mutant and tested reporter expression at the restrictive temperature 15°C. We confirmed that at 15°C *nhr-67(ot202*ts) worms lack expression of the GABAergic reporter *unc-25p::GFP* in RME neurons. However, the RMEs affected were RMEL/R (**fig. S3, A** and **B**), not RMED/V, which are the neurons affected in cohesin mutants (**Fig. 3, A** and **B**). Furthermore, approximately 11% of *nhr-67(ot202*ts) worms had extra tyraminergic-like cells, but those cells similarly were not RMED/V (**fig. S3, C** and **D**). We also examined *ceh-10*, another known regulator of RME cell identity (*32*). *ceh-10(gm127)* mutant worms exhibit defective expression of GABAergic reporters in RMED (*32*). We used CRISPR to create the *ceh-10(n6695)* allele, which is identical to *ceh-10(gm127)*. We confirmed that *ceh-10(n6695)* mutants have reduced expression of *unc-25p::GFP* in RMED (**fig. S3, E** and **F**), but these mutants did not generate extra RIM-like cells (**fig. S3, G**), showing that *ceh-10* does not inhibit expression of a tyraminergic-like fate by RMED. Together, these data indicate that neither *nhr-67* nor *ceh-10* acts in the cohesin pathway that inhibits fate transformation from GABAergic RMED/V to tyraminergic-like neuronal fates.

### The PLZF transcription factor EOR-1 acts with cohesin to prevent GABAergic neurons from acquiring a tyraminergic identity

Our EMS mutagenesis screens also identified three mutations (*n5163*, *n5294* and *n6588*) that cause extra *tdc-1p::4xNLS::GFP*-positive cells but were not alleles of *coh-1* (**fig. S4A**). We found that *n5163*, *n5294* and *n6588* mutants contain opal, amber and amber premature stop codons, respectively, in the gene *eor-2* (**fig. S4B**). We confirmed that these *eor-2* mutations were responsible for the generation of extra GFP-positive cells by testing another *eor-2* allele, *cs42*, and by rescuing the *eor-2* isolates (*n5163*, *n5294* and *n6588*) with wild-type copies of *eor-2* (**fig. S4C**). *eor-2* encodes a protein that acts as a cofactor of the transcription factor EOR-1, a homolog of the mammalian promyelocytic leukemia zinc finger (PLZF) transcription factor (*34–36*). We showed that *eor-1(cs28)* null mutants (*36*) also generate extra adrenergic-like cells (**fig. S4, D** and **E**). Furthermore, *eor-1(cs28)* and *eor-2(n5163)* mutations did not show additive effects in the *eor-1(cs28); eor-2(n5163)* double mutant (**fig. S4, D** and **E**), suggesting that *eor-1* and *eor-2* act together to prevent the generation of extra adrenergic-like cells.

We next asked if the extra adrenergic-like cells were extra tyraminergic-like or extra octopaminergic-like cells. Similar to cohesin mutants, *eor-1* mutants generated both extra tyraminergic-like and extra octopaminergic-like cells, and the generation of extra tyraminergic-like and octopaminergic-like cells was rescued by wild-type copies of *eor-1* fused with a *gfp* gene (*eor-1::GFP*) (**Fig. 4, A** and **B**, and **fig. S4, F** and **G**). Because the positions of the extra tyraminergic-like cells in *eor-1* mutants were similar to those in cohesin mutants, we asked if the extra cells expressing the tyraminergic reporter were the GABAergic RME neurons by examining the expression of the tyraminergic GFP reporter and the GABAergic *unc-47::SL2::H2B::mChOpti* reporter in *eor-1* mutants as we did for *coh-1* mutants. Indeed, the extra tyraminergic-like cells were the GABAergic RMED, RMEV or RMEL neurons (**Fig. 4, C** and **D**), indicating that *eor-1* inhibits the expression of a tyraminergic fate by the GABAergic RME neurons, as does cohesin.

**Fig. 4.**
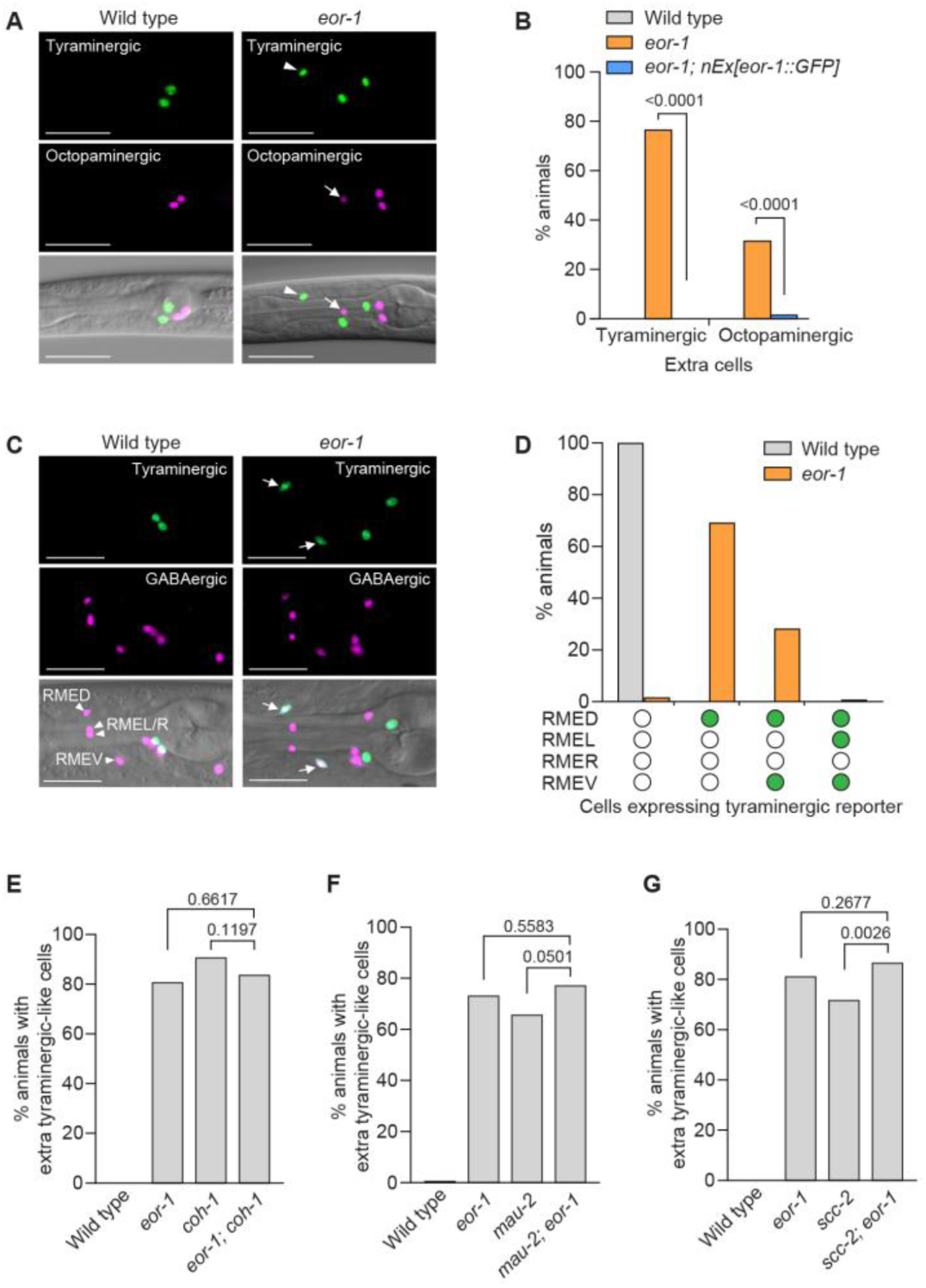
Cohesin and EOR-1 act in the same pathway to inhibit the neuronal GABAergic-to-adrenergic fate transformation. (**A**) The extra tyraminergic-like and octopaminergic-like cells in *eor-1(cs28)* mutants are different cells. Strains expressed an *F23H12.7p::4xNLS::GFP* (tyraminergic) and a *tbh-1p::4xNLS::mCherry* (octopaminergic) reporter. Arrowheads, extra tyraminergic-like cells. Arrows, extra octopaminergic-like cells. Scale bar, 20 µm. Worms at the L2-L4 larval stages were analyzed in panels **A**, **B** and **G**. (**B**) Percentages of wild-type and *eor-1(cs28)* mutant animals displaying extra tyraminergic-like and octopaminergic-like cells. Wild-type copies of *eor-1* fused with a GFP sequence at the 3’ end (*nEx3176[eor-1::GFP]*) rescue the generation of extra tyraminergic-like and octopaminergic-like cells. p-value, Fisher’s exact test. n=60 animals. (**C**) The extra tyraminergic-like cells are the GABAergic RMED and RMEV neurons in *eor-1(cs28)* mutants. Strains expressed an *F23H12.7p::4xNLS::GFP* (tyraminergic) and an *unc-47::SL2::H2B::mChOpti* (GABAergic) reporter in panels **C** and **D**. Arrowheads, GABAergic RMED, RMEL, RMER and RMEV. Arrows, extra tyraminergic-like cells. Scale bar, 20 µm. Worms at the L4 larval stage were analyzed in panels **C** and **D**. (**D**) Quantification of RME neurons expressing the tyraminergic reporter in *eor-1(cs28)* mutants. Empty circle, RME with no GFP expression. Green circle, RME with GFP expression. n=120 animals. (**E** to **G**) Percentages of animals expressing extra tyraminergic-like cells in wild-type animals, *eor-1(cs28)* and *coh-1(n6618)* single mutants, *eor-1(cs28); coh-1(n6618)* double mutants (**E**), *mau-2(qm160)* single mutants and *mau-2(qm160); eor-1(cs28)* double mutants (**F**), and *scc-2(n6656)* single mutants and *scc-2(n6656); eor-1(cs28)* double mutants (**G**). p-value, Fisher’s exact test. n=120 animals for wild type, *eor-1* and *coh-1*, and 117 for *eor-1; coh-1* (**E**). n=120 animals for wild type, *eor-1* and *mau-2*, and 132 for *mau-2; eor-1* (**F**). n=160 animals for wild type, *eor-1* and *scc-2*, and 135 for *scc-2; eor-1* (**G**). Strains expressed an *F23H12.7p::4xNLS::GFP* (tyraminergic) reporter. Worms at the L1-L2 larval stages were analyzed for panels **E** and **F**.

Given the similar phenotypes of cohesin and *eor-1* mutants, we hypothesized that cohesin and *eor-1* act together. Consistent with this hypothesis, a previous study reported that *eor-1* and the cohesin gene *mau-2* act in the same pathway during the post-embryonic maturation of the *C. elegans* HSN neurons (*37*). To determine if cohesin and *eor-1* act in the same pathway to inhibit expression of a tyraminergic fate by the normally GABAergic RMEDs and RMEVs, we generated cohesin and *eor-1* double mutants. *eor-1; coh-1* and *mau-2; eor-1* double mutants reached fertile adulthood, but their progeny arrested as larvae, consistent with the previously reported maternal effects of *eor-1* and *mau-2* mutations (*37*, *38*). We assayed the number of extra tyraminergic-like cells in the arrested larvae and found that the *coh-1* and *mau-2* mutations did not enhance the number of extra tyraminergic-like cells in *eor-1* mutants (**Fig. 4, E** and **F**). We next tested the cohesin gene *scc-2.* The progeny of *scc-2; eor-1* double mutants grew to adulthood and produced viable offspring. Like *coh-1* and *mau-2* mutations, an *scc-2* mutation did not enhance the generation of the extra tyraminergic-like cells in *eor-1* mutants (**Fig. 4G**). These data are consistent with the hypothesis that cohesin and *eor-1* act in the same pathway to prevent the generation of tyraminergic-like RMED/V cells. Furthermore, similar to cohesin mutants, *eor-1* mutants displayed defects in the expression of the GABA biosynthesis reporter *unc-25p::GFP* (*18*, *39*, *40*) and of the RMED/V reporter *ins-13p::GFP* and, like cohesin mutants, had truncated RMED and RMEV neuronal processes (**fig. S4, H** to **K**). Taken together, these data indicate that cohesin and *eor-1* likely act together to inhibit the expression of a tyraminergic fate by the normally GABAergic RMED and RMEV neurons.

### The NuRD complex and TRA-4 (another PLZF homolog) promote a tyraminergic fate in the absence of cohesin or *eor-1* function

Cohesin-mediated changes in genomic architecture are known to influence the activities of diverse chromatin remodeling complexes and transcription factors (*9*). To identify chromatin factors that act in the cohesin pathway, we performed an RNAi suppressor screen using *coh-1* mutants that ectopically express the tyraminergic GFP reporter *F23H12.7p::4xNLS::GFP* (**Fig. 5A**). Specifically, we tested available RNAi clones against 374 chromatin-modifying and - related genes (*41*) and found that knockdown of either *lin-40* or *dcp-66* (mammalian counterparts MTA and p66), which encode components of the nucleosome remodeling and deacetylase (NuRD) complex, suppressed the generation of extra tyraminergic-like cells in *coh-1* mutants (**Fig. 5, B** and **C**). The NuRD complex is a major evolutionarily conserved chromatin remodeler that modulates chromatin state and gene expression via histone deacetylation (*42–46*). RNAi against *lin-40* and *dcp-66* similarly suppressed the generation of extra tyraminergic-like cells in other cohesin (*mau-2*, *scc-2*) and *eor-1* mutants (**Fig. 5C**). In addition, RNAi against other components of the NuRD complex – *mbd-2*, *let-418*, *lin-53* and *hda-1* (mammalian counterparts MBD, Mi-2, RbAp and HDAC) – also suppressed the generation of extra tyraminergic-like cells in cohesin and *eor-1* mutants (**Fig. 5, D** and **E**). These data indicate that the NuRD complex is required for the misexpression of a tyraminergic fate in cohesin and *eor-1* mutants.

**Fig. 5.**
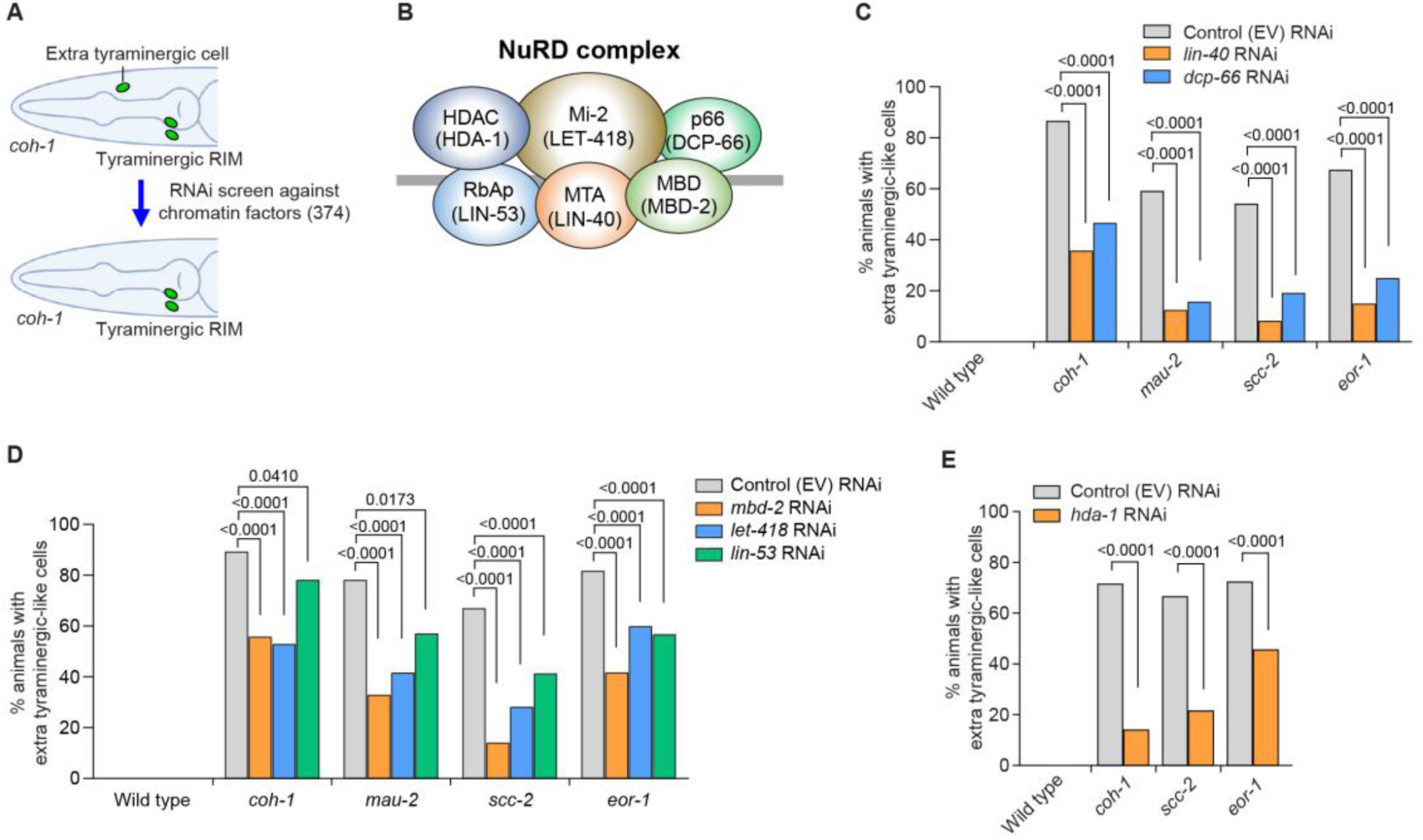
The NuRD complex promotes an alternative fate by RME neurons in the absence of cohesin or EOR-1. (**A**) Schematic showing a candidate suppressor RNAi screen against chromatin factors to identify genes required for the generation of extra tyraminergic-like cells in *coh-1(n6618)* mutants. Green ovals indicate expression of a tyraminergic RIM GFP reporter (*F23H12.7p::4xNLS::GFP*). (**B**) Components in the nucleosome remodeling and deacetylase (NuRD) complex. *C. elegans* orthologs are shown in parentheses. (**C**) *lin-40* RNAi and *dcp-66* RNAi partially suppress the generation of extra tyraminergic-like cells in *coh-1(n6618)*, *mau-2(qm160)*, *scc-2(n6656)* and *eor-1(cs28)* mutants. *F23H12.7p::4xNLS::GFP* was used as a tyraminergic reporter in panels **C** to **E**. EV, empty vector. p-value, Fisher’s exact test. n=120 animals. Worms at the L2-L4 larval stages were analyzed for panels **C** to **E**. (**D**) *mbd-2* RNAi, *let-418* RNAi and *lin-53* RNAi partially suppress the generation of extra tyraminergic-like cells in *coh-1(n6618)*, *mau-2(qm160)*, *scc-2(n6656)* and *eor-1(cs28)* mutants. EV, empty vector. p-value, Fisher’s exact test. n=170 animals for control RNAi, *mbd-2* RNAi and *let-418* RNAi in all five genotypes. n=108 animals for *lin-53* RNAi in wild type, 55 animals for *lin-53* RNAi in *coh-1*, 35 animals for *lin-53* RNAi in *mau-2*, 67 animals for *lin-53* RNAi in *scc-2(n6656)*, and 132 animals for *lin-53* RNAi in *eor-1*. Smaller numbers of *lin-53* RNAi-treated animals were scored, because *lin-53* RNAi caused incompletely penetrant lethality. (**E**) Knockdown of *hda-1* by RNAi partially suppresses the generation of extra tyraminergic-like cells in *coh-1(n6618)*, *scc-2(n6656)* and *eor-1(cs28)* mutants. EV, empty vector. p-value, Fisher’s exact test. n=120 animals.

In addition to EOR-1, the *C. elegans* genome encodes a second PLZF ortholog, TRA-4 (**Fig. 6A**). A previous study found that TRA-4 interacts with the histone deacetylase HDA-1, a NuRD complex component, in sex determination (*47*). To test the possibility that TRA-4 acts with the NuRD complex to mediate the neuronal fate transformation in cohesin and *eor-1* mutants, we examined the effect of a putative null allele of *tra-4(bc250)* (*47*) in cohesin and *eor-1* mutant backgrounds. Interestingly, the *tra-4* mutation suppressed the generation of extra tyraminergic-like cells in cohesin and *eor-1* mutants; this effect was not additive in *eor-1; hda-1; tra-4* triple mutants (**Fig. 6, B** and **C**, and **fig. S5, A** and **B**). These data indicate that *tra-4* and *hda-1* act in the same pathway. We confirmed that expression of wild-type copies of *tra-4* fused with GFP (*GFP::tra-4*) rescued the effect of *tra-4* mutation in *eor-1; tra-4* double mutants, showing loss of *tra-4* function suppresses the generation of extra tyraminergic-like cells in *eor-1* mutants (**Fig. 6D**). Furthermore, a *tra-4* mutation significantly suppressed the reduced expression of *unc-25p::GFP* and *ins-13p::GFP* in cohesin and *eor-1* mutants (**Fig. 5, E** and **F**, and **fig. S5, C** to **F**). By contrast, *tra-4* or *hda-1* mutations did not affect the expression of a tyraminergic reporter in the RIMs of worms that were wild-type for cohesin and *eor-1*, indicating that neither the NuRD complex nor TRA-4 is necessary for the expression of normal tyraminergic fates (**fig. S5, G** and **H**). These data indicate that EOR-1 and TRA-4, the two *C. elegans* orthologs of mammalian PLZF, play opposite roles in the fate determination of the GABAergic RMED and RMEV neurons.

**Fig. 6.**
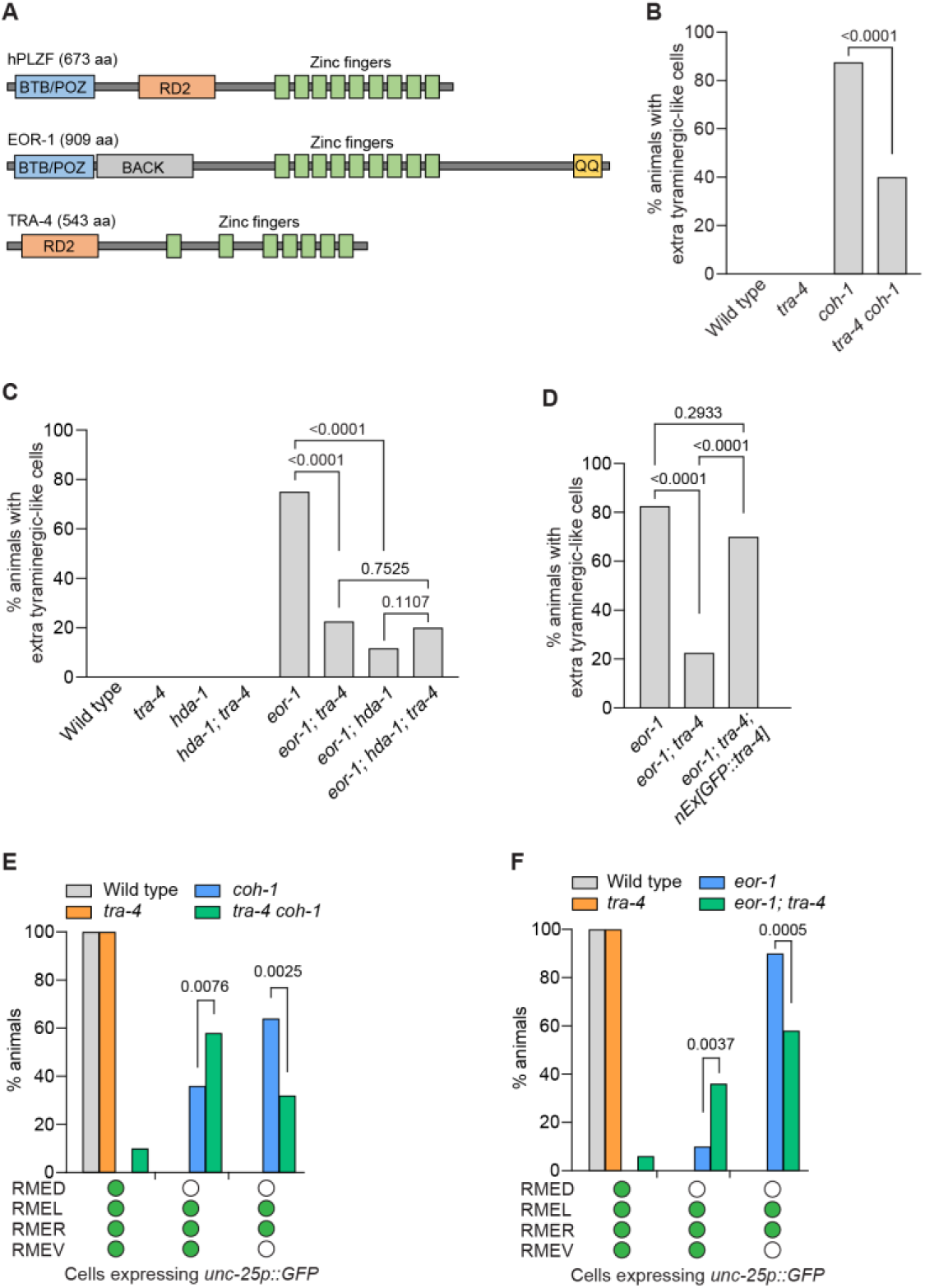
TRA-4 acts with the NuRD complex to promote an alternative fate by RME neurons in the absence of cohesin or EOR-1. (**A**) Simplified domain structures of human PLZF (hPLZF), EOR-1 and TRA-4. BTB/POZ, bric a brac, tramtrack, broad complex/poxvirus and zinc finger. RD2, second repression domain. BACK, BTB and C-terminal Kelch. QQ, polyglutamine. aa, amino acid. Adapted from Hoeppner et al. (2004); Grote and Conradt. (2006); Howell et al. (2010) (*36*, *47*, *68*). Worms at the L2-L4 larval stages were analyzed in panels **B** to **F**. (**B**) The *tra-4(bc250)* mutation partially suppresses the generation of extra tyraminergic-like cells in *coh-1(n6618)* mutants. p-value, Fisher’s exact test. n=120 animals. (**C**) *tra-4(bc250)* and *hda-1(cw2)* suppress the generation of extra tyraminergic-like cells in *eor-1(cs28)* mutants, and the effects of *tra-4(bc250)* and *hda-1(cw2)* mutations are not additive in *eor-1(cs28); hda-1(cw2); tra-4(bc250)* triple mutants. p-value, Fisher’s exact test. n=120 animals. (**D**) Extrachromosomal copies of wild-type *tra-4* fused with GFP at the 5’ end (*nEx3242[GFP::tra-4]*) rescue the effect of *tra-4(bc250)* for the generation of extra tyraminergic-like cells in *eor-1(cs28); tra-4(bc250)* double mutants. p-value, Fisher’s exact test. n=40 animals. (**E** and **F**) The *tra-4(bc250)* mutation partially suppresses the defects in the expression of *unc-25p::GFP* (GABAergic reporter) in the RME neurons in *coh-1(n6618)* mutants (**E**) and *eor-1(cs28)* mutants (**F**). Empty circle, RME with no GFP expression. Green circle, RME with GFP expression. p-value, Fisher’s exact test. n=50 animals.

### The two PLZF-like proteins play opposite roles in controlling expression of genes regulated by cohesion

That cohesin and *eor-1* mutants share the same phenotype and that EOR-1 is a transcription factor led us to hypothesize that cohesin and EOR-1 work together to regulate the expression of the same target genes and that *tra-4* antagonizes the expression of those targets. To test this hypothesis, we performed RNA-Seq using wild-type, *coh-1* and *eor-1* single-mutant, and *tra-4 coh-1* and *eor-1; tra-4* double-mutant embryos. Principal component analysis (PCA) showed that *coh-1* mutants have a greater effect on gene expression than *eor-1* mutants (**Fig. 7A**). Changes in overall gene expression in *coh-1* and *eor-1* mutants displayed significantly positive correlation (r=0.89, p-value <0.001) (**Fig. 7B**). Differentially expressed gene analyses (DEG, adjusted p-value <0.05 and fold change >2) showed that *coh-1* mutants have approximately 4-fold more DEGs than *eor-1* mutants (**Fig. 7C** and **fig. S6A**). Although overall changes of gene expression were strongly correlated between *coh-1* and *eor-1* mutant embryos, the p-values and fold changes of genes changed in *eor-1* mutants were below our DEG criteria (adjusted p-value <0.05 and fold change >2). These values are consistent with our observation that although *coh-1* and *eor-1* mutants are similar in phenotype, the *coh-1* mutant phenotype is stronger than the *eor-1* mutant phenotype. Strikingly, most genes (97%) down-regulated in *eor-1* mutant embryos but only 24% of genes down-regulated in *coh-1* mutant embryos were down-regulated in both *coh-1* and *eor-1* mutants, indicating that cohesin has broader activities in gene expression and that EOR-1 acts on a subset of COH-1 target genes (**Fig. 7C**). Gene enrichment analysis using WormCat (*48*) showed that the genes down-regulated in both *coh-1* and *eor-1* mutants are involved in mRNA function, cell cycle and development, which are all processes potentially associated with neuronal development (**Fig. 7D**). However, because the mutations analyzed are likely pleiotropic and because the enrichment categories identified by WormCat were based on data generated from whole embryos, these categories might reflect broad transcriptional changes in whole embryos rather than specific pathway regulation in specific cell types. Similarly, 82% of genes up-regulated in *eor-1* mutant embryos, and 20% of genes up-regulated in *coh-1* mutant embryos were up-regulated in both *coh-1* and *eor-1* mutants (**fig. S6A**). The genes up-regulated in both *coh-1* and *eor-1* mutants did not yield any significantly enriched WormCat categories. Together, these data suggest that cohesin regulates many of the transcriptional targets regulated by EOR-1.

**Fig. 7.**
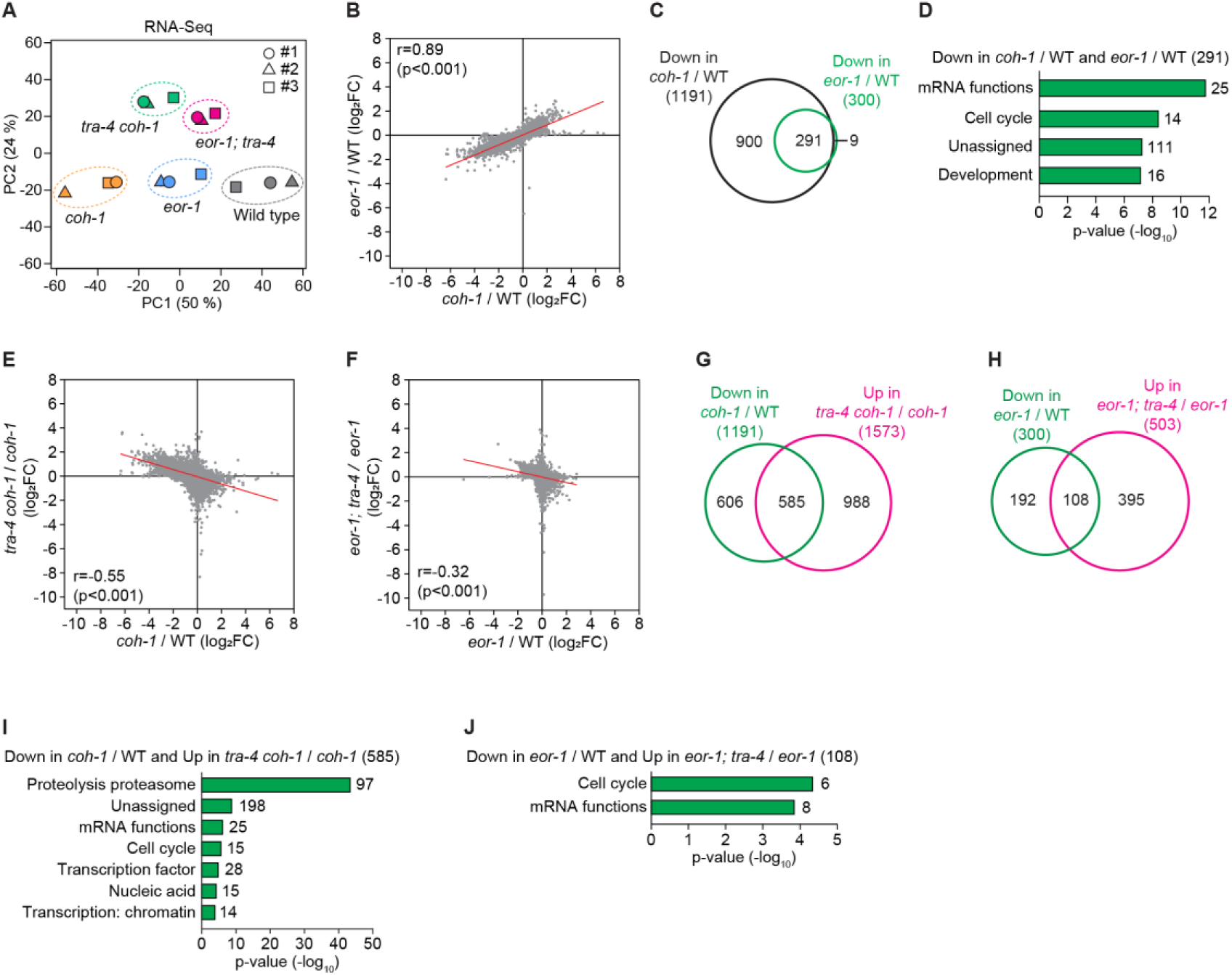
TRA-4 antagonizes the expression of transcriptional targets of cohesin and EOR-1. (**A**) Principal component analysis (PCA) plots showing the variations in RNA-Seq data obtained from wild-type, *coh-1(n6618)*, *eor-1(cs28)*, *tra-4(bc250) coh-1(n6618)*, and *eor-1(cs28); tra-4(bc250)* embryos in three replicates (#1, #2, #3). (**B**) Dot plot showing the correlation of fold changes (log_2_FC) of mRNA expression between *coh-1(n6618)* and *eor-1(cs28)* mutants versus wild-type (WT) embryos. Red line, simple linear regression. r and p-value, Spearman correlation. (**C**) Venn diagram showing overlap between genes down-regulated in *coh-1(n6618)* and *eor-1(cs28)* mutants. Criteria for differentially expressed genes (DEGs): adjusted p-value <0.05 and fold change >2. The DEGs are shown in **Supplementary Data S1**. (**D**) Gene set enrichment terms of the 291 genes down-regulated in both *coh-1(n6618)* and *eor-1(cs28)* mutant embryos. The gene set enrichment was analyzed using WormCat (*48*), which applies multiple error corrections – Fisher’s exact test to calculate the enrichment and p-value and Bonferroni correction for the represented categories. Only categories meeting the Bonferroni-corrected threshold (< 0.05) are shown in panels **D**, **I** and **J**. (**E** and **F**) Dot plots showing a negative correlation of fold changes (log_2_FC) of mRNA expression in *tra-4(bc250) coh-1(n6618)* double-mutant versus *coh-1(n6618)* single-mutant embryos (**E**), and *eor-1(cs28); tra-4(bc250)* double-mutant versus *eor-1(cs28)* single-mutant embryos (**F**). Red line, simple linear regression. r and p-value, Spearman correlation. (**G** and **H**) Venn diagrams showing overlap between genes down-regulated in *coh-1(n6618)* versus wild-type embryos and genes up-regulated in *tra-4(bc250) coh-1(n6618)* double-mutant versus *coh-1(n6618)* single-mutant embryos (**G**), and genes down-regulated in *eor-1(cs28)* single-mutant versus wild-type and genes up-regulated in *eor-1(cs28); tra-4(bc250)* double-mutant versus *eor-1(cs28)* single-mutant embryos (**H**). Adjusted p-value <0.05 was used as a criterion for DEGs. The DEGs are shown in **Supplementary Data S1**. (**I** and **J**) Gene set enrichment results of the overlapping genes in panels **G** (**I**) and **H** (**J**).

Next we asked if *tra-4* regulates the expression of targets of cohesin and EOR-1. By comparing gene expression changes in *tra-4 coh-1* double mutants versus *coh-1* single mutants and *eor-1; tra-4* double mutants versus *eor-1* single mutants, we showed that the global changes in gene expression in *coh-1* single mutants were reversed in *tra-4 coh-1* double mutants (r=-0.55, p-value <0.001), and a similar reverse correlation was observed between *eor-1* single mutants and *eor-1; tra-4* double mutants (r=-0.32, p-value <0.001) (**Fig. 7, E** and **F**). Furthermore, DEG (adjusted p-value <0.05) analyses showed that almost half (49%) of the genes down-regulated in *coh-1* were significantly up-regulated in *tra-4 coh-1* double mutants, and 36% of genes down-regulated in *eor-1* single mutants were significantly up-regulated in *eor-1; tra-4* double mutants (**Fig. 7, G** and **H**). The *tra-4* mutation caused a lesser effect on the genes up-regulated in *coh-1* or *eor-1* mutants than on the genes down-regulated in *coh-1* or *eor-1* mutants (**fig. S6, B** and **C**). Gene set enrichment analysis showed that the *tra-4*-suppressed genes were enriched in gene expression and development-related terms, such as mRNA function and cell cycle (**Fig. 7, I** and **J**). Collectively, these data suggest that TRA-4 acts downstream of (or parallel to) cohesin and EOR-1 to antagonize the expression of transcriptional targets regulated by cohesin and EOR-1.

## Discussion

The nervous system consists of diverse types of neurons, and genetic and epigenetic factors drive distinct neuronal fates. In this study we demonstrate that cohesin, the nucleosome remodeling and deacetylase (NuRD) complex and two promyelocytic leukemia zinc finger (PLZF) transcription factors interact in the determination of specific neuronal fates in *C. elegans* (**Fig. 8**). By analyzing mutants isolated from EMS mutagenesis screens using a GFP reporter expressed in tyraminergic and octopaminergic cells, we found that cohesin and *eor-1*, which encodes a homolog of the mammalian PLZF transcription factor, act in the same genetic pathway to prevent a subset of GABAergic neurons from expressing genes normally expressed in tyraminergic cells. An RNAi suppressor screen revealed that the NuRD complex facilitates the fate change from GABAergic to tyraminergic that occurs in the absence of cohesin or *eor-1* function. TRA-4, another PLZF homolog, acts with the NuRD complex. The NuRD complex functions in cell-fate determination in multiple tissues in *C. elegans,* including the somatic gonad, germline, and intestinal and hypodermal cells, as well as in the expression of apoptotic cell death (*41*, *43–46*, *49–57*). However, no previous study has described a role for the NuRD complex in determining specific neuronal fates. Our data show that cohesin functions with the EOR-1 transcription factor to drive the GABAergic identity of specific neurons by inhibiting expression of genes characteristic of a tyraminergic neuronal fate, and that the NuRD complex and TRA-4 promote the tyraminergic fate in a subset of GABAergic neurons when cohesin or EOR-1 is not functional. We suggest that during evolution the NuRD complex and TRA-4 were required for the expression of an adrenergic fate by primordial RMED/V neurons and that the emergence or activation of cohesin and EOR-1 inhibited these primordial factors, thereby facilitating the origin of GABAergic RMED/V neurons.

**Fig. 8.**
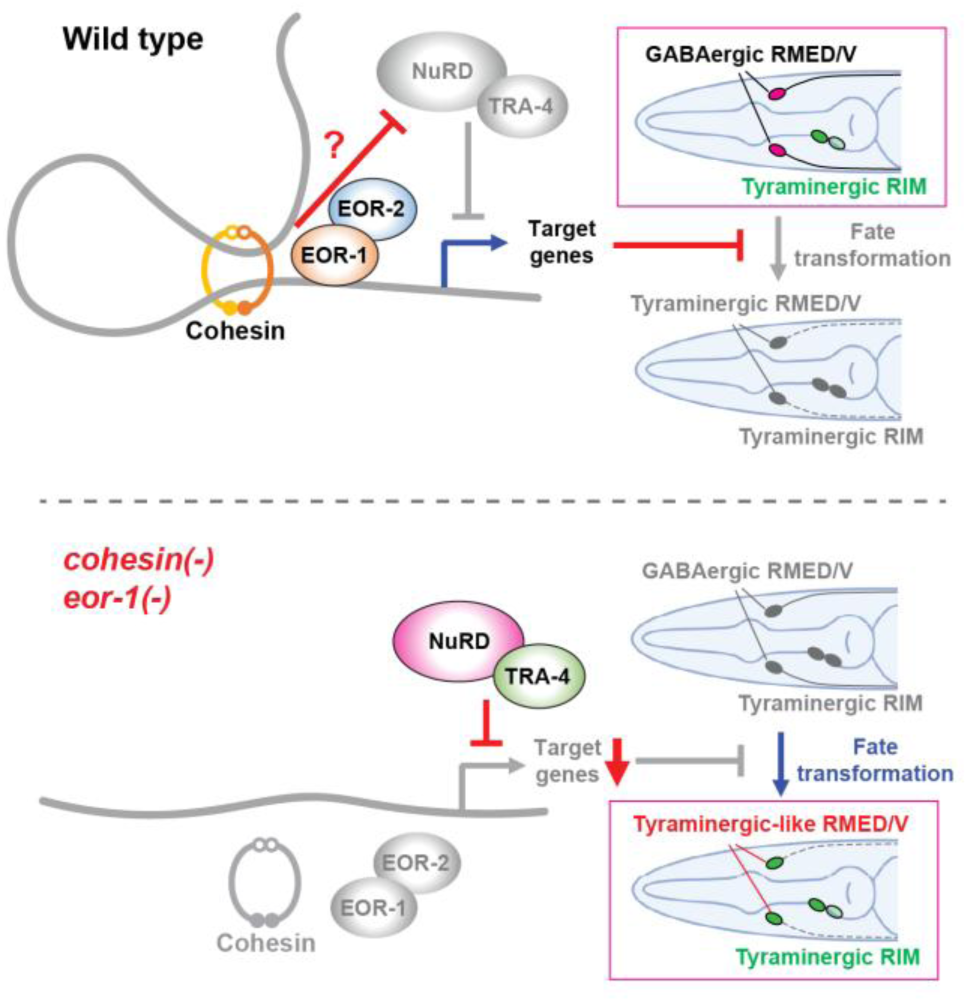
Cohesin and EOR-1 drive specific neuronal fates by inhibiting fate transformation from GABAergic to tyraminergic. Model showing cohesin, the EOR-1 transcription factor and the EOR-2 co-factor acting together to prevent a cell-fate transformation of RMED and RMEV from GABAergic to tyraminergic-like through transcriptional regulation. We speculate that there is an inhibitory action of cohesin or EOR-1 on the NuRD complex and TRA-4 in wild-type animals. In cohesin mutants, the fate of the normally GABAergic RMED and RMEV neurons is transformed into a tyraminergic-like fate. The NuRD complex and TRA-4 facilitate this cell-fate transformation by repressing a subset of the targets of cohesin and EOR-1.

Since cohesin modulates genomic architecture and EOR-1 regulates transcription, the spatiotemporal control of their DNA binding is likely crucial for the functional interaction between them. Although our genetic and transcriptomic data demonstrate that cohesin and EOR-1 act in the same pathway, how they interact is unclear. We suggest three possible models. First, EOR-1 might alter chromatin structure and recruit cohesin to specific genomic regions. Second, cohesin might guide EOR-1 to specific genomic regions by modulating chromatin structure. Third, other factors might independently recruit both cohesin and EOR-1 to their target regions. A previous report showed that in *C. elegans* EOR-1 can genetically interact with the chromatin-remodeling complexes SWI/SNF (switching and sucrose fermentation) and RSC (remodeling the structure of chromatin) (*37*, *58*). Also, EOR-1-binding sites are enriched in less accessible chromatin regions in early embryonic stages (*59*). These findings suggest that EOR-1 might act as a pioneer factor that binds to closed chromatin regions to make those regions accessible. Interestingly, two recent papers show that cohesin can be associated with DNA loops extruded from potentially active enhancer regions (*60*, *61*), consistent with a model in which EOR-1 recruits cohesin to potential enhancer regions to promote gene expression by mediating the enhancer-promoter interaction.

We demonstrate that cohesin and EOR-1, a *C. elegans* ortholog of mammalian PLZF, function together to promote a specific cellular fate by inhibiting an alternative fate. Accumulating evidence suggests that PLZF proteins have similar cell-fate inhibitory roles across species. For example, the PLZF ortholog Tramtrack promotes glial cell development by inhibiting neuronal differentiation in Drosophila (*62–64*). *tramtrack* mutant embryos display supernumerary neuronal cells (*65*), and *tramtrack* is both necessary and sufficient to prevent fate transformation from support cells to neurons (*66*). These studies concluded that PLZF likely determines cellular fates between glial or support cells and differentiated neurons by inhibiting neuronal differentiation. Similarly, in chicken and mouse embryos, PLZF inhibits premature differentiation of neurons and maintains glial progenitors (*67*). Our findings raise the intriguing possibility that cohesin cooperates with PLZF to regulate development across species.

The *C. elegans* genome encodes two presumptive PLZF homologs, EOR-1 and TRA-4. These two proteins are known to play diverse roles in tissue development, neuronal maturation, programmed cell death and sex determination (*18*, *34–37*, *39*, *47*, *68*), but had not been previously shown to act together, let alone act together in an antagonistic manner. The human PLZF protein contains three functional domains: BTB/POZ (bric a brac, tramtrack, broad complex/poxvirus and zinc finger), RD2 (second repression domain) and nine zinc fingers (*69*). EOR-1 and TRA-4, the two *C. elegans* PLZF orthologs, have different domains conserved with PLZF: EOR-1 shares a BTB/POZ domain and nine zinc fingers, while TRA-4 shares an RD2 domain and seven zinc fingers. The conserved zinc-finger domains suggest that EOR-1 and TRA-4 might bind to the same DNA sequences. Additionally, TRA-4 interacts with HDA-1, a component of the generally repressive NuRD complex (*47*), and we found that TRA-4 antagonizes the transcriptional function of EOR-1, suggesting that TRA-4 acts as a transcriptional repressor. Interestingly, a number of ZBTB proteins, which contain BTB/POZ and zinc finger domains, have been shown to interact with cohesin, and depletion of ZBTB7B and ZBTB21 decreases the chromatin binding of cohesin (*70*). This result is consistent with our model that EOR-1, which contains a BTB/POZ domain, directly or indirectly interacts with cohesin to change the local genomic architecture. Since the orthologous domains of human PLZF are separated in the two *C. elegans* PLZF orthologs, in future studies it might be feasible to disentangle the roles of human PLZF domains by analyzing the distinct functions of the two *C. elegans* orthologs in controlling gene expression and genome organization.

Previous ChIP-Seq and ATAC-Seq data showed that chromatin-binding of EOR-1 and HDA-1, a component of the NuRD complex, exhibit opposite patterns as worms develop from early embryonic to larval stages: the EOR-1-binding motif is enriched in less accessible chromatin regions in early embryonic stages compared to late larval (L3) stages, while the HDA-1-binding motif is enriched in open chromatin regions in early embryonic stages compared to late larval (L3) stages (*71*). This finding is consistent with our data showing that EOR-1 and the NuRD complex function oppositely in the control of the RMED/V GABAergic neuronal fates. Our genetic data suggest that the NuRD complex acts downstream of (or parallel to) EOR-1, but whether EOR-1 and cohesin affect NuRD complex activity and chromatin state is still unknown.

Although we analyzed fate changes of specific neurons, our transcriptomic analysis of whole-embryo data does not reveal the precise cellular contexts in which cohesin and EOR-1 function. It will be important to identify the specific genomic loci at which cohesin and EOR-1 act. Future studies using approaches such as single-cell RNA-seq, single-cell CUT&RUN and ChIP-Seq, and spatial transcriptomics will be needed to determine the functions of cohesin and EOR-1 and their genomic targets at single-cell resolution.

Cornelia de Lange syndrome (CdLS) is one of the best-studied human disorders associated with cohesin dysfunction (*72*). Notably, cells from individuals with CdLS, as well as Nipbl (an ortholog of *C. elegans scc-2*)-deficient mice, Drosophila and zebrafish models with reduced Nipbl or cohesin dosage, exhibit widespread alterations in gene expression (*73–76*). These findings are consistent with our observation that cohesin, PLZF and the NuRD complex function as key regulators of transcriptional programs. To our knowledge, cohesin, PLZF transcription factors, and the NuRD complex have never before been linked in a shared molecular genetic pathway. The observation that inhibition of the NuRD complex can ameliorate defects in cohesin and *eor-1* mutants suggests potential therapeutic approaches to genetic disorders caused by mutations in the cohesin or PLZF genes. Such strategies might hold promise not only for neurodevelopmental syndromes but also for hematologic malignancies and solid tumors in which this pathway is disrupted (*69*, *72*).

## Materials and Methods

### Strains

*C. elegans* strains were maintained on standard nematode growth media (NGM, 3 g/liter NaCl, 2.5 g/liter peptone and 17 g/liter agar supplemented with 1 mM CaCl_2_, 1 mM MgSO_4_, 1 mM KPO_4_ and 5 mg/liter cholesterol) with *E. coli* OP50 as a food source (*77*). All *C. elegans* strains were maintained at 20°C except for the temperature-sensitive *lin-15AB(n765ts)* and *pha-1(e2123ts)* strains, which were maintained at 15°C. A complete list of strains used in this study is presented in **Supplementary Data S2**.

### Molecular cloning and strain construct

Molecular cloning was performed using the In-Fusion cloning kit (Takara Bio) or the method of bacterial homologous recombination (*78*). Germline transformation was performed by injecting plasmids into the gonads in day-1 adult worms (*79*). CRISPR strains were created as described previously (*80–82*). Integration of extrachromosomal arrays was performed as described previously (*83*) and outcrossed at least three times. To construct *tbh-1p::4xNLS::mCherry* plasmid, the promoter region of *tbh-1* (3.4 kb upstream from the start codon) was amplified by PCR and inserted into the plasmid containing *4xNLS::mCherry::unc-54 3’UTR*. The putative promoter of *F23H12.7* (2.2 kb) was amplified by PCR and inserted into the plasmid containing *4xNLS::mCherry::unc-54 3’UTR* or *4xNLS::GFP::unc-54 3’UTR*. *scc-2(n6656)* was created by knocking out 2,960 bp using CRISPR/Cas9. *scc-2(n6656)* is an in-frame deletion spanning from half of the 1st exon to a portion of the 7th exon of the *scc-2* gene. To create *coh-1p::coh-1::linker::wrmScarlet::3xFLAG::coh-1 3’UTR* plasmid, the *coh-1* promoter (4.2 kb), coding region and 3’UTR were PCR-amplified and cloned into the pPD95.75 backbone without *unc-54* 3’UTR, and then flexible linker 3x(Gly-Gly-Ser-Gly), wrmScarlet and 3xFLAG sequences were inserted between the *coh-1* coding region and the *coh-1* 3’UTR. PCR-amplified *eor-2* promoter (2 kb), coding region and 3’UTR were purified using the QIAquick PCR Purification Kit (QIAGEN), and injected into *eor-2* isolates (*n5163*, *n5294* and *n6588*) for rescue assays. To construct *eor-1p::eor-1::linker::GFP::HA::eor-1 3’UTR*, the promoter (1.2 kb), coding region and 3’UTR of *eor-1* were amplified by PCR and inserted into the pPD95.75 backbone without *unc-54* 3’UTR, and 3x(Gly-Gly-Ser-Gly), GFP and HA sequences were inserted between the coding region and the *eor-1* 3’UTR. To create *tra-4p::3xFLAG::GFP::linker::tra-4::tra-4 3’UTR* plasmid, the 1 kb promoter, coding region and 3’UTR of *tra-4* were amplified and inserted into the pPD95.75 backbone without *unc-54* 3’UTR, and the 3xFLAG, GFP and 3x(Gly-Gly-Ser-Gly) were inserted between the promoter and coding region. A *tra-4* transgene with the linker::GFP sequence at the 3’ end did not rescue *tra-4(bc250)* mutants, suggesting that the GFP tag at the C-terminus disrupts the function of TRA-4. To create *ins-13p::GFP::3xFLAG::unc-54 3’UTR* plasmid, 2 kb of the putative *ins-13* promoter was amplified by PCR and inserted into the plasmid containing *GFP::3xFLAG::unc-54 3’UTR*. *nEx2914*, *nEx3036*, *nEx3072*, *nEx3060* and *nEx3210* were used to create *nIs874*, *nIs878*, *nIs883* and *nIs884*, *nIs889* and *nIs896* by UV-integration, respectively. The plasmid and primer sequences used in this study are available upon request. Injected plasmid concentrations are described in **Supplementary Data S2**.

### EMS mutagenesis screens

Ethyl methanesulfonate (EMS) mutagenesis screens were performed as described previously with some modifications (*84*). *nIs180[tdc-1p::4xNLS::GFP]* or *nIs180; rol-6(su1006); nIs349[ceh-28p::4xNLS::mCherry]* worms at the L4 stage were treated with 47 mM EMS (Sigma) in M9 buffer for 4 hrs at room temperature with constant rotation. The *rol-6(su1006)* mutation was used for visualizing the laterally positioned RIML/R and RICL/R neurons in rolling worms, and the M4 neuron-specific marker *nIs349* was added to exclude potential mutations in the canonical cell-death genes *egl-1*, *ced-9*, *ced-4* and *ced-3*. We used two screen designs, first with only *nIs180[tdc-1p::4xNLS::GFP]* and second with *nIs180; rol-6; nIs349*. Mutagenized F2 worms were screened for extra and reduced number of GFP-positive cells using a Nikon SMZ18 fluorescence dissecting microscope. Genomic DNA was prepared using the Puregene Cell and Tissue Kit (Qiagen, #158388), and library preparation and whole-genome sequencing were performed in the BioMicro Genomic Center at MIT. Whole-genome sequencing data were analyzed using the Mutation Identification in Model Organism Genomes using Desktop PCs (MiModD) pipeline on the Galaxy platform (*85*). The Hawaiian variant mapping and EMS density mapping methods were used to identify mutations (*86*). Isolated mutants were crossed to the Hawaiian strain CB4856 for the Hawaiian variant mapping (*87*).

### Fluorescence imaging

Worms at the L2-L4 stages were anesthetized with 10-30 mM sodium azide (Sigma) in M9 buffer and mounted on 2-5% agarose pads for imaging. Embryos were transferred into M9 buffer and mounted on 2-5% agarose pads. Epifluorescence micrographs were taken using a Zeiss Axio Imager Z2 widefield fluorescence microscope. Multiple images were merged to show cells in different focal planes if necessary. A Zeiss LSM800 was used to capture confocal micrographs. Confocal Z-stacks were merged to show cells in different focal planes. Image processing was performed using Microsoft PowerPoint, ZEN 2.5 blue edition (ZEISS) and Fiji (ImageJ) (*88*).

### Quantification of worms with extra cells

Worms at the L2-L4 larval stages were used for quantification. The numbers of worms expressing extra GFP-positive and mCherry-positive cells on NGM plates were scored using a Nikon SMZ18 fluorescence dissecting microscope. p-values were calculated by Fisher’s exact test using the GraphPad Prism software. To count the number of or identify fluorescence-positive cells in single worms, anesthetized worms mounted on 2-5% agarose pads were analyzed using a Zeiss Axio Imager Z2 widefield fluorescence microscope.

### Quantification of the length of RMED and RMEV processes

Worms at the L4 larval stage were anesthetized with 10-30 mM sodium azide (Sigma) in M9 buffer and mounted on 2-5% agarose pads for imaging. The *ins-13p::GFP* signals in the neuronal processes of RMED and RMEV were captured using a Zeiss Axio Imager Z2 widefield fluorescence microscope. The lengths of neuronal processes were measured using Fiji (ImageJ) (*88*). p-values were calculated by one-way ANOVA using the GraphPad Prism software.

### Embryo isolation by bleaching

Worms were cultivated in 100 mm high growth (HG) NGM plates (*89*) seeded with 1 ml *E. coli* NA22 cultured in LB. For an initial synchronization, gravid adult worms and larvae were collected and washed using 10 ml M9 buffer by centrifugation at 180 x g for 1 min multiple times until the supernatant was clear. The worm pellet was resuspended in 7 ml M9 buffer, and 0.7 ml NaOH (5 M) and 0.7 ml bleach (Clorox, 7.55% sodium hypochlorite) were added to release embryos. When most of the worms was lysed (approximately 5-7 min depending on the number of worms), released embryos were washed with 10 ml M9 buffer three times by centrifugation at 180 x g for 1 min. The washed embryos were incubated in 1 ml M9 buffer at 20°C overnight with constant nutating. The number of arrested L1 worms was estimated by counting worms in 1 µl M9 buffer. Because *eor-1* and *coh-1* mutants produce fewer eggs than the wild type, approximately 10,000 wild-type, 15,000 *eor-1* mutant and 20,000 *coh-1* mutant L1 worms were transferred onto each HG plate seeded with NA22. Synchronized L1 worms grown on HG plates were incubated at 20°C for three days. Synchronized adult worms were bleached again as described above to obtain a large population of embryos.

### RNA isolation and RNA-Seq

Wild-type, *coh-1(n6618)*, *eor-1(cs28)*, *tra-4(bc250) coh-1(n6618)* and *eor-1(cs28); tra-4(bc250)* strains expressing *nIs883* (tyraminergic GFP/octopaminergic mCherry reporter) were used for RNA-Seq. Three biological replicates were prepared independently on the same days. Embryos were released by bleaching adult worms in 2-3 HG plates seeded with NA22 per condition as described above. The released embryos were incubated in M9 buffer at 20°C by shaking until most embryos developed to the comma to 2-fold stages. Embryos were collected by centrifugation at 180 x g for 1 min, and the supernatant was discarded. Embryos in 100 µl M9 buffer were stored at -80°C. Total RNA was isolated using the RNeasy kit (QIAGEN). Embryos in M9 buffer were resuspended in 800 µl of RLT buffer (QIAGEN RNeasy kit) containing 1% β-mercaptoethanol (Mallinckrodt) and dissociated using a BeadBug microtube homogenizer (Benchmark Scientific) with zirconium beads (0.5 mm, Benchmark Scientific) for 30 sec. Dissociated embryos and beads were centrifuged at 15,800 x g for 10 min at room temperature to separate debris. The supernatant was used to isolate total RNA according to the manufacturer’s instructions. Sequencing libraries were prepared by the MIT BioMicro Center, and sequencing was performed using the NEXTSeq500 sequencing platform with a 75 nt setup. Differentially expressed genes were analyzed using WBcel235 as a reference genome, STAR (v2.5.3) (*90*), RSEM (v1.3.1) (*91*) and DESeq2 (v1.24.0) (*92*). The raw data are available in Gene Expression Omnibus (GEO): GSE283114. RPKM (reads per kilobase per million mapped reads) values were used for PCA (principal component analysis). Venn diagrams were generated using DeepVenn (https://www.deepvenn.com/). Gene set enrichment analysis was performed using WormCat (*48*).

### RNA interference (RNAi) experiments

Gene-specific double-stranded RNA-expressing *E. coli* (HT115) was cultured in liquid LB media containing ampicillin (75 µg/ml) at 37°C overnight. RNAi bacteria were seeded onto NGM plates containing ampicillin (75 µg/ml) and IPTG (isopropyl β-D-1-thiogalactopyranoside, 1 mM, Amresco) and incubated at 37°C for 24 hr. L3-L4 larvae were transferred onto RNAi bacteria-seeded NGM plates, and phenotypes were scored at the next generation. The sequences of RNAi clones targeting cohesin subunits and the NuRD complex used in this study were confirmed by sequencing. The chromatin factor RNAi clones used for the suppressor screen were cultured from the Ahringer RNAi library (*93*). Individual RNAi clones targeting *smc-3*, *him-1*, *dcp-66*, *lin-40*, *let-418* and *mbd-2* were cultured from the Ahringer RNAi library, and *smc-3*, *hda-1* and *lin-53* were cultured from the Vidal RNAi library (*94*).

## Supporting information

Lee et al., Supplementary Data S1_RNA-Seq

Lee et al., Supplementary Data S2_Strains

## Acknowledgments

We thank members of the Horvitz laboratory for helpful discussions.

## Author contributions

Investigation: DL, TH

Conceptualization, methodology, resources, project administration, writing – original manuscript, and writing – review and editing: HRH, DL

Data curation, validation, formal analysis, software, and visualization: DL Funding acquisition and supervision: HRH

## Data, code, and materials availability

The raw RNA-Seq data are available in Gene Expression Omnibus (GEO): GSE283114. All data are available in the main text or the supplementary materials. This study did not generate new code or new materials.

## Supplementary Materials

Supplementary Materials: Figures S1-S6

Supplementary Data S1: Excel file containing RNA-Seq data

Supplementary Data S2: Excel file containing strain and allele list

