## Supplementary Figures and Legends for "Cohesin and NuRD Antagonistically Drive Alternative Neuronal Fates via PLZF Transcription Factors"

Dongyeop Lee *et al*.

**This PDF file includes:**

Figs. S1 to S6

**Other Supplementary Materials for this manuscript include the following:**

Supplementary Data S1 and S2

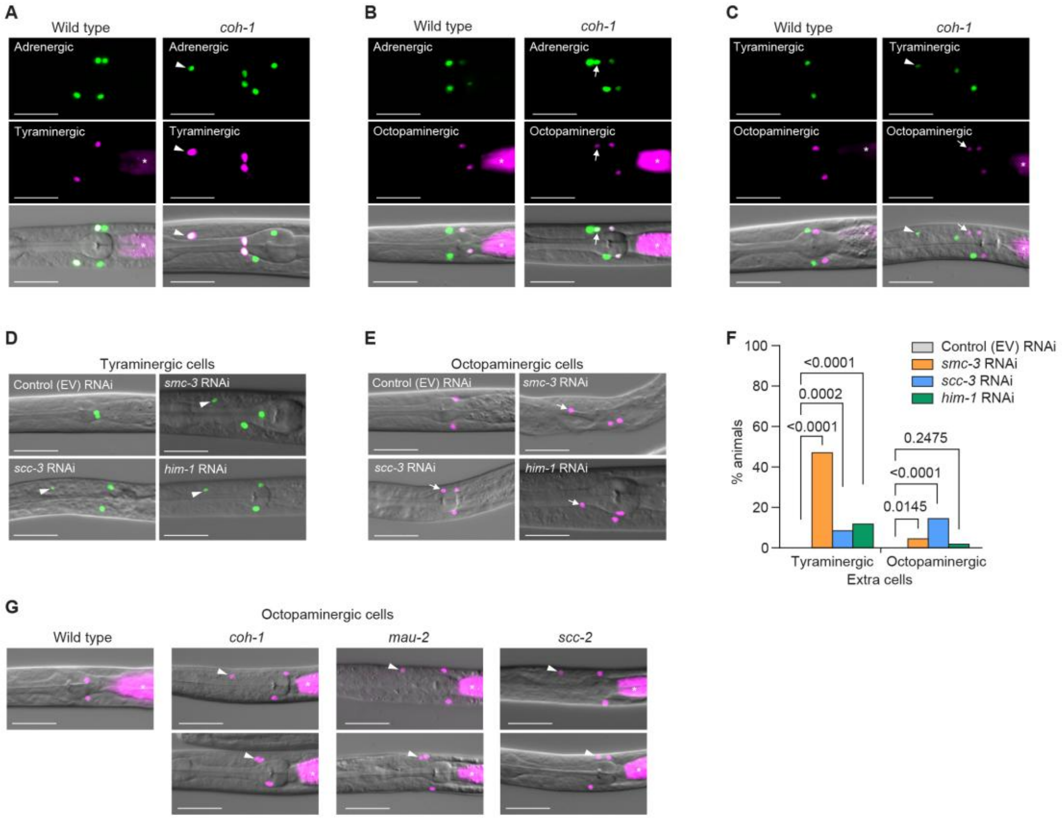


**Fig. S1. Genetic inhibition of cohesin components generates extra tyraminergic-like and extra octopaminergic-like cells.** (**A** and **B**) *coh-1(n6618)* mutants generate both extra tyraminergic-like (**A**) and octopaminergic-like cells (**B**). All worms carried a *tdc-1p::4xNLS::GFP* (adrenergic) and either an *F23H12.7p::4xNLS::mCherry* (tyraminergic) or a *tbh-1p::4xNLS::mCherry* (octopaminergic) reporter. Arrowheads, extra tyraminergic-like cells. Arrows, extra octopaminergic-like cells. Asterisks, co-injection marker (*ges-1p::mCherry*). Scale bar, 20 µm. Worms at the L2-L4 larval stages were analyzed for panels **A** to **G**. (**C**) The extra tyraminergic-like and octopaminergic-like cells in *coh-1(n6618)* mutants are different cells. All animals carried both an *F23H12.7p::4xNLS::GFP* (tyraminergic) and a *tbh-1p::4xNLS::mCherry* (octopaminergic) reporter. Arrowheads, extra tyraminergic-like cells. Arrows, extra octopaminergic-like cells. Asterisks, co-injection marker (*ges-1p::mCherry*). Scale bar, 20 µm. (**D** and **E**) RNAi against *smc-3*, *scc-3* and *him-1* generates extra tyraminergic-like cells (**D**) and octopaminergic-like cells (**E**). Arrowheads, extra tyraminergic-like cells. Arrows, extra octopaminergic-like cells. EV, empty vector. Scale bar, 20 µm. (**F**) Quantification of panels **D** and **E**. Percentages of worms exhibiting extra tyraminergic-like and extra octopaminergic-like cells are shown. EV, empty vector. p-value, Fisher's exact test. n=150 animals. (**G**) The positions of extra octopaminergic-like cells in *coh-1(n6618)*, *mau-2(qm160)* and *scc-2(n6656)* mutants are variable, suggesting different sources of extra octopaminergic-like cells. Strains contained a *tbh-1p::4xNLS::GFP* (octopaminergic) reporter. Arrowheads, extra octopaminergic-like cells. Asterisks, co-injection marker (*ges-1p::mCherry*). Scale bar, 20 µm.

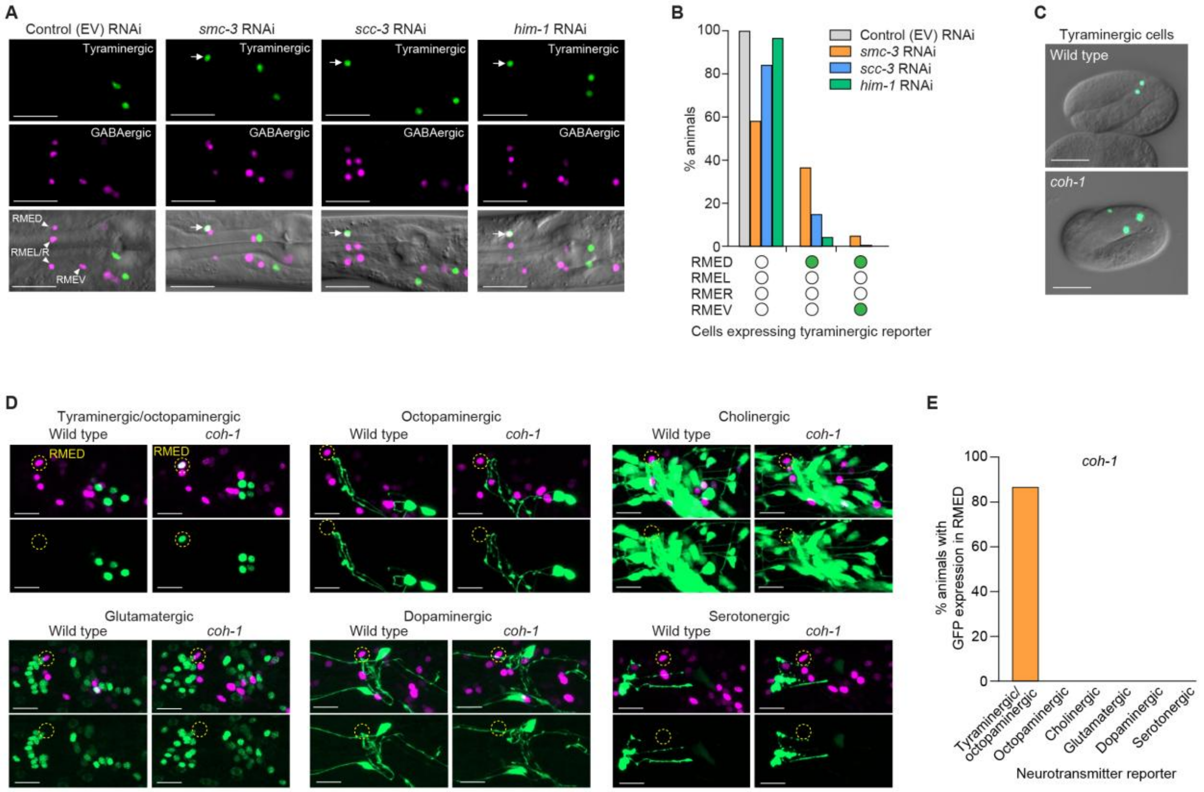


**Fig. S2. Cohesin prevents a tyraminergic fate for RMED and RMEV.** (**A**) The extra tyraminergic-like cell in *smc-3*, *scc-3* and *him-1* RNAi-treated worms is RMED. Animals expressed a *F23H12.7p::4xNLS::GFP* (tyraminergic) and an *unc-47::SL2::H2B::mChOpti* (GABAergic) reporter in panels **A** and **B**. Arrowheads, GABAergic RMED, RMEL, RMER and RMEV. Arrows, extra tyraminergic-like cells. Scale bar, 20 µm. EV, empty vector. Worms at the L4 larval stage were analyzed for panels **A**, **B**, **D** and **E**. (**B**) Percentage of animals expressing the tyraminergic reporter in RMED and/or RMEV. Empty circle, RME with no GFP expression. Green circle, RME with GFP expression EV, empty vector. n=120 animals. (**C**) Representative images showing the expression of *F23H12.7p::4xNLS::GFP* (tyraminergic) in wild-type and *coh-1(n6618)* mutant embryos (three-fold stage). The *coh-1(n6618)* mutant embryo displays an extra tyraminergic-like cell. Scale bar, 20 µm. (**D**) Confocal fluorescence micrographs showing the expression of *unc-47::SL2::H2B::mChOpti* (GABAergic) and neurotransmitter GFP reporters in wild-type and *coh-1* mutant animals. The *tdc-1p::4xNLS::GFP* (tyraminergic/octopaminergic) reporter is expressed in RMED, while other neurotransmitter reporters such as *tbh-1p::GFP* (octopaminergic), *unc-17/cha-1p::GFP* (cholinergic), *eat-4::SL2::YFP::H2B* (glutamatergic), *dat-1::T2A::NeonGreen* (dopaminergic) or *tph-1p::GFP* (serotonergic) are not expressed in RMED. Upper panels in each reporter show merged GFP/mChOpti images and lower panels show GFP images. *coh-1(n6618)* was present in all strains except those carrying the cholinergic GFP reporter. *coh-1(n6718)*, a mutation identical to *n6618*, was created using CRISPR/Cas9 in the cholinergic GFP reporter background, because the cholinergic reporter and the *coh-1* gene are in the same chromosome and creating the identical *coh-1* mutation by CRISPR/Cas9 was more efficient than crossing the reporter to *coh-1* mutants. Dashed circles, RMED. Scale bar, 10 µm. (**E**) The proportion of animals with GFP expression in RMED. n=38 animals for tyraminergic/octopaminergic, 45 animals for octopaminergic, 41 animals for cholinergic, 42 animals for glutamatergic, 50 animals for dopaminergic, and 44 animals for serotonergic reporters.

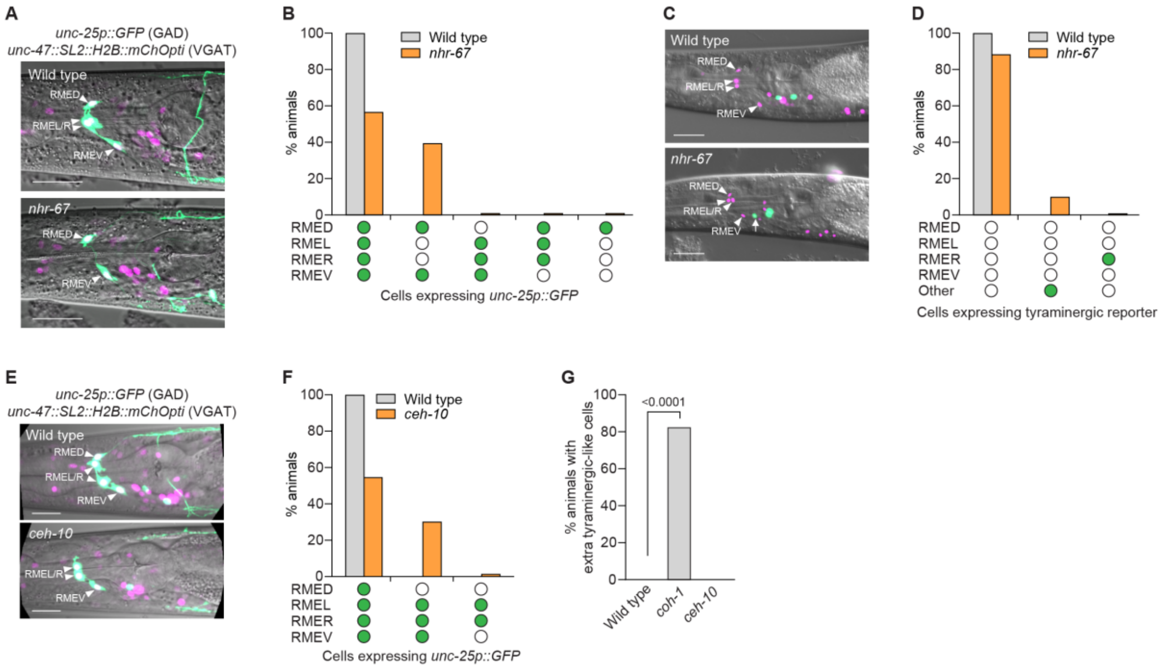


**Fig. S3. Neither *nhr-67* nor *ceh-10* act in the cohesin pathway.** (**A**) Confocal fluorescence micrographs showing that GAD is not expressed in RMEL or RMER in *nhr-67(ot202*ts*)* mutants. Animals that expressed the reporters *unc-25p::GFP* (GAD) and *unc-47::SL2::H2B::mChOpti* (VGAT) are shown in **A** and **B**. Arrowheads, RME neurons expressing GFP. Scale bar, 20 µm. Worms were grown at 15°C for panels **A** to **D**. L4 worms were analyzed for panels **A** to **G**. (**B**) Percentages of wild-type animals and *nhr-67(ot202*ts*)* mutants that express *unc-25p::GFP* (GAD) in RMEs. Empty circle, RME with no GFP expression. Green circle, RME with GFP expression. n=120 animals. (**C**) The majority of extra tyraminergic-like cells in *nhr-67(ot202)* mutants are not RMEs. Animals that expressed the reporters *F23H12.7p::4xNLS::GFP* (tyraminergic) and *unc-47::SL2::H2B::mChOpti* (GABAergic) are shown in panels **C** and **D**. Arrowheads, GABAergic RMED, RMEL, RMER and RMEV. Arrow, extra tyraminergic-like cell. Scale bar, 20 µm. (**D**) Percentages of wild-type animals and *nhr-67(ot202*ts*)* mutants that have cells expressing the tyraminergic-like reporter *F23H12.7p::4xNLS::GFP*. Empty circle, cells with no GFP expression. Green circle, cells with GFP expression. n=90 animals. (**E**) Confocal fluorescence micrographs showing that GAD is not expressed in RMED in *ceh-10(n6695)* mutants. Strains that expressed the reporters *unc-25p::GFP* (GAD) and *unc-47::SL2::H2B::mChOpti* (VGAT) are shown in panels **E** and **F**. Arrowheads, RME neurons expressing GFP. Scale bar, 20 μm. (**F**) Percentages of wild-type animals and *ceh-10(n6695)* mutants that express *unc-25p::GFP* (GAD) in RMEs. Empty circle, RME with no GFP expression. Green circle, RME with GFP expression. n=120 animals. (**G**) Percentage of wild-type, *coh-1(n6618)* and *ceh-10(n6695)* mutant animals displaying extra tyraminergic-like cells. *ceh-10(n6695)* mutants do not generate extra tyraminergic-like cells. Animals expressed the *F23H12.7p::4xNLS::GFP* (tyraminergic) and *unc-47::SL2::H2B::mChOpti* (GABAergic) reporters.

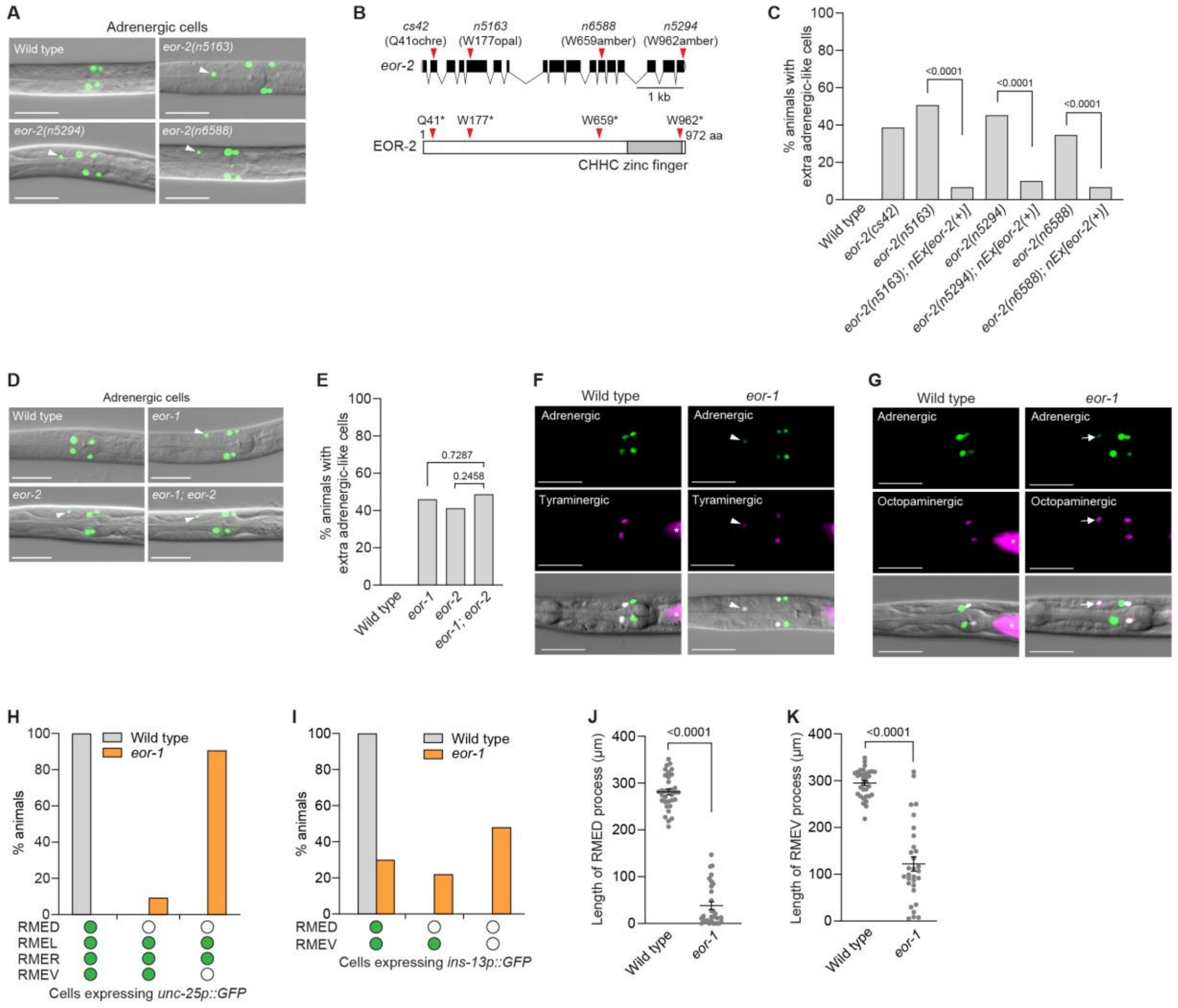


**Fig. S4. EOR-1 inhibits generation of extra tyraminergic-like cells and promotes a GABAergic fate in RMED/V.** (**A**) Three *eor-2* alleles (*n5163*, *n5292* and *n6588*) obtained from EMS mutagenesis screens generate extra adrenergic-like cells. Animals expressed a *tdc-1p::4xNLS::GFP* (adrenergic) reporter in panels **A**, **C**, **D** and **E**. Arrowheads, extra tyraminergic-like cells. Scale bar, 20 µm. Worms at the L2-L4 larval stages were analyzed in panels **A**, and **C** to **I**. (**B**) The *eor-2(n5163)*, *eor-2(n5294)* and *eor-2(n6588)* mutations isolated in this study and another *eor-2(cs42)* allele generate opal, amber, amber and ochre premature stop codons in the gene *eor-2*, respectively. The CHHC zinc finger domain is highlighted in grey. (**C**) Percentages of wild-type and *eor-2* mutant animals displaying extra adrenergic-like cells, and the effect of rescue with extrachromosomal copies of wild-type *eor-2*. *nEx3030[eor-2(+)]*, *nEx3031[eor-2(+)]* and *nEx3051[eor-2(+)]* rescues *eor-2(n5163)*, *eor-2(n5294)* and *eor-2(n6588)*, respectively. p-value, Fisher's exact test. n=150 animals. (**D**) *eor-1(cs28)*, *eor-2(n5163)* and *eor-1(cs28); eor-2(n5163)* double mutants generate extra adrenergic-like cells. Arrowheads, extra adrenergic-like cells. Scale bar, 20 µm. (**E**) The effects of *eor-1(cs28)* and *eor-2(n5163)* single mutants are not additive in *eor-1(cs28); eor-2(n5163)* double mutants. p-value, Fisher's exact test. n=150 animals. (**F** and **G**) *eor-1(cs28)* mutants generate extra tyraminergic-like (**F**) and extra octopaminergic-like cells (**G**). Strains contained a *tdc-1p::4xNLS::GFP* (adrenergic) and either an *F23H12.7p::4xNLS::mCherry* (tyraminergic) or a *tbh-1p::4xNLS::mCherry* (octopaminergic) reporter. Arrowheads, extra tyraminergic-like cells. Arrows, extra octopaminergic-like cells. Asterisk, co-injection marker (*ges-1p::mCherry*). Scale bar, 20 µm. (**H** and **I**) Percentages of wild-type and *eor-1(cs28)* mutant animals that express *unc-25p::GFP* (**H**) and *ins-13p::GFP* (**I**) in RMEs. Empty circle, RME with no GFP expression. Green circle, RME with GFP expression. n=150 animals. (**J** and **K**) Quantification of the length of RMED (**J**) and RMEV (**K**) in wild-type animals and *eor-1(cs28)* mutants. Error bars, mean ± SEM. p-value, one-way ANOVA. n=33 animals for wild type and 30 animals for *eor-1(cs28)*. Worms at the L4 larval stage were analyzed. Please note that the same wild-type conditions were used in **Fig. 3, F** and **G**.

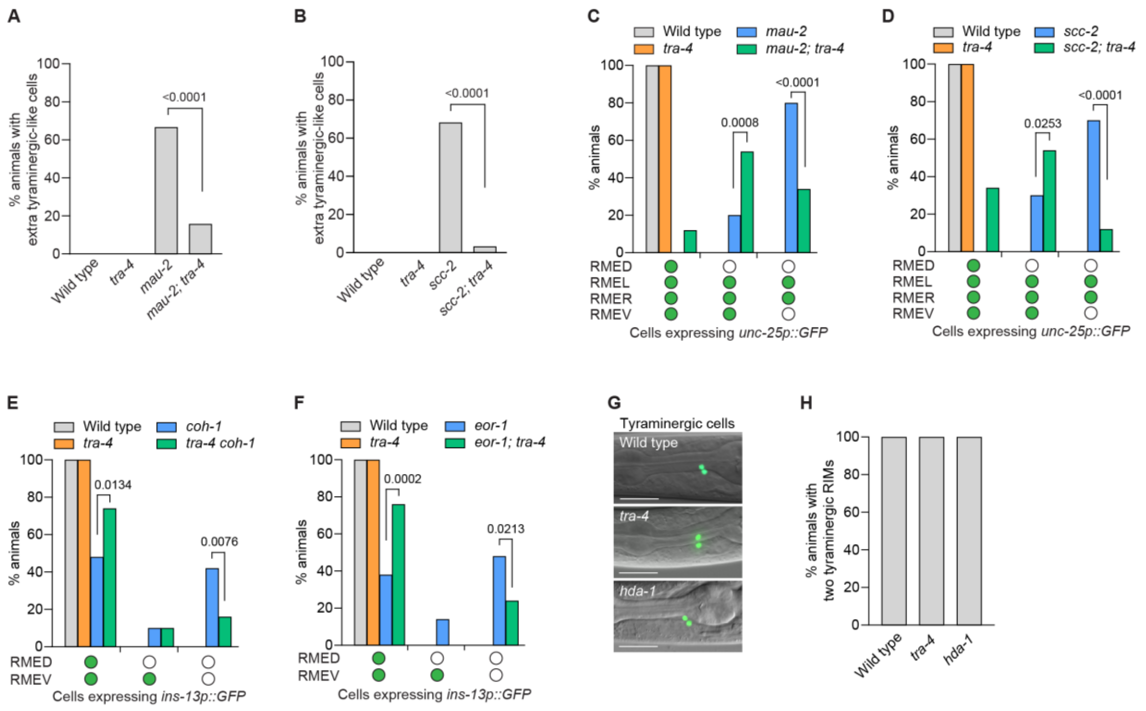


**Fig. S5. TRA-4 promotes an alternative fate by RME neurons in cohesin and *eor-1* mutants.** (**A** and **B**) The *tra-4(bc250)* mutation partially suppresses the generation of extra tyraminergic-like cells in *mau-2(qm160)* mutants (**A**) and *scc-2(n6656)* mutants (**B**). All animals expressed the tyraminergic reporter *F23H12.7p::4xNLS::GFP*. p-value, Fisher's exact test. n=120 animals. Worms at the L2-L4 larval stages were analyzed. (**C** and **D**) The *tra-4(bc250)* mutation partially suppresses the defects in the expression of *unc-25p::GFP* (GABAergic reporter) in the RME neurons in *mau-2(qm160)* mutants (**C**) and *scc-2(n6656)* mutants (**D**). Empty circle, RME with no GFP expression. Green circle, RME with GFP expression. p-value, Fisher's exact test. n=50 animals. Worms at the L4 larval stage were analyzed. (**E** and **F**) The *tra-4(bc250)* mutation partially suppresses the defects in the expression of *ins-13p::GFP* (RMED/V reporter) in RMED and/or RMEV in *coh-1(n6618)* mutants (**E**) and *eor-1(cs28)* mutants (**F**). Empty circle, RME with no GFP expression. Green circle, RME with GFP expression. p-value, Fisher's exact test. n=50 animals. Worms at the L2-L4 larval stages were analyzed. (**G** and **H**) *F23H12.7p::4xNLS::GFP* (tyraminergic reporter) expression at the L4 larval stage (**G**) and the percentages of animals with two tyraminergic RIM neurons (**H**) in the heads of wild-type animals, and *tra-4(bc250)* and *hda-1(cw2)* mutants. Scale bar, 20 µm. n=120 animals. Worms at the L2-L4 larval stages were analyzed in panel **H**.

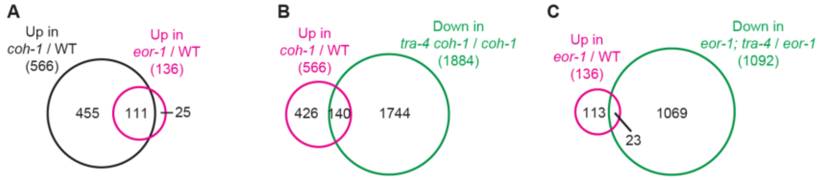


**Fig. S6. *tra-4* mutation has marginal effects on genes up-regulated in *coh-1* and *eor-1* mutant embryos.** (**A**) Venn diagram showing overlap between genes up-regulated in *coh-1(n6618)* and *eor-1(cs28)* mutant embryos. Adjusted p-value <0.05 and fold change >2 were used as criteria for differentially expressed genes. (**B** and **C**) Venn diagrams showing overlaps between genes up-regulated in *coh-1(n6618)* mutant compared to wild-type embryos and down-regulated in *tra-4(bc250) coh-1(n6618)* double-mutant compared to *coh-1(n6618)* single-mutant embryos (**B**), and genes up-regulated in *eor-1(cs28)* mutant compared to wild-type embryos and genes down-regulated in *eor-1(cs28); tra-4(bc250)* double-mutant compared to *eor-1(cs28)* single-mutant embryos (**C**).

**Supplementary Data S1. (separate file)**

Excel file containing RNA-Seq data

**Supplementary Data S2. (separate file)**

Excel file containing strains and alleles used in this study
